# Selective functional inhibition of a tumor-derived p53 mutant by cytosolic chaperones identified using split-YFP in budding yeast

**DOI:** 10.1101/2021.04.02.438278

**Authors:** Ashley S. Denney, Andrew D. Weems, Michael A. McMurray

**Affiliations:** Department of Cell and Developmental Biology, University of Colorado Anschutz Medical Campus, Aurora, CO 80045

**Keywords:** p53, septin, mutant, protein folding, molecular chaperone, heat shock protein 90 (Hsp90), bimolecular fluorescence complementation (BiFC), *Saccharomyces cerevisiae*

## Abstract

Life requires the oligomerization of individual proteins into higher-order assemblies. In order to form functional oligomers, monomers must adopt appropriate three-dimensional structures. Molecular chaperones transiently bind nascent or misfolded proteins to promote proper folding. Single missense mutations frequently cause disease by perturbing folding despite chaperone engagement. A misfolded mutant capable of oligomerizing with wild-type proteins can dominantly poison oligomer function. We previously found evidence that human-disease-linked mutations in *Saccharomyces cerevisiae* septin proteins slow folding and attract chaperones, resulting in a kinetic delay in oligomerization that prevents the mutant from interfering with wild-type function. Here we build upon our septin studies to develop a new approach to identifying chaperone interactions in living cells, and use it to expand our understanding of chaperone involvement, kinetic folding delays, and oligomerization in the recessive behavior of tumor-derived mutants of the tumor suppressor p53. We find evidence of increased binding of several cytosolic chaperones to a recessive, misfolding-prone mutant, p53(V272M). Similar to our septin results, chaperone overexpression inhibits the function of p53(V272M) with minimal effect on the wild type. Unlike mutant septins, p53(V272M) is not kinetically delayed under conditions in which it is functional. Instead, it interacts with wild-type p53 but this interaction is temperature sensitive. At high temperatures or upon chaperone overexpression, p53(V272M) is excluded from the nucleus and cannot function or perturb wild-type function. Chaperone inhibition liberates the mutant to enter the nucleus where it has a slight dominant-negative effect. These findings provide new insights into the effects of missense mutations.

## INTRODUCTION

Higher-order protein structures are essential for life. Protein monomers must be synthesized, correctly folded, and efficiently assembled into homo- or hetero-oligomers while avoiding inappropriate interaction with misfolded partners. Quality control at each step is important to ensure these processes are accurate and timely. Molecular chaperones aid in quality control by assisting protein folding through repetitive cycles of client binding and release. A survey of nearly 3,000 human disease-causing missense mutant proteins revealed that ∼30% showed increased association with chaperones relative to the wild-type (WT) allele, and this value is likely an underestimate (Sahni *et al*. 2015). This finding illustrates an important principle: although chaperone binding to WT clients promotes client folding to the native state, in the case of misfolded mutants chaperone engagement does not guarantee that the client will be released in a form that is competent for function. While sequestration of grossly misfolded proteins by chaperones is an established mechanism for maintaining cellular fitness (Escusa-Toret *et al*. 2013), the idea that persistent chaperone engagement may actually interfere with normal protein function has received much less attention.

Septins are highly conserved cytoskeletal proteins whose assembly into hetero-oligomers is important for various cellular processes (Mostowy and Cossart 2012; Woods and Gladfelter 2021). Septins were originally discovered in *Saccharomyces cerevisiae* as temperature-sensitive (TS) mutants that failed to complete cytokinesis at high temperature (Hartwell 1971). We previously demonstrated that at low temperatures permissive for cellular function many TS septin mutants are kinetically delayed in their folding, and that this delay has consequences for oligomerization when mutant septin monomers compete with WT monomers (Johnson *et al*. 2015), a quality control phenomenon we termed kinetic allele discrimination in oligomerization (KADO) (Schaefer *et al*. 2016). Exclusion of the mutant septin from septin hetero-oligomers ensures that the mutant allele is recessive and does not interfere with the function of the WT allele (Johnson *et al*. 2015). We noticed that the excluded mutant persisted in the cytosol complexed with non-septin proteins, and identified these as cytosolic chaperones (Johnson *et al*. 2015). Deletion of genes encoding some of the same cytosolic chaperones perturbed KADO and allowed TS-mutant septins to incorporate into hetero-oligomers despite the presence of the WT (Johnson *et al*. 2015). Thus chaperone binding to the mutant septin contributed to the kinetic competition between the two alleles. In the case of a mutant that is non-functional at all temperatures, chaperone deletion converted the otherwise recessive mutant septin to a dominant-lethal allele, presumably by allowing the mutant to incorporate into septin hetero-oligomers and “poison” septin function (Johnson *et al*. 2015). Finally, chaperone overexpression inhibited septin function in cells in which only the mutant septin allele was expressed, consistent with a model in which chaperone engagement sequesters the mutant protein and blocks its ability to assemble into functional hetero-oligomers (Johnson *et al*. 2015). One yeast septin mutant subject to KADO carries the same amino acid substitution as a human mutation that blocks septin hetero-oligomerization and prevents septin function in sperm (Kuo *et al*. 2012; Weems *et al*. 2014; Johnson *et al*. 2015). We wanted to understand more about which chaperones bind WT and mutant septins and the extent to which the mechanisms we discovered with mutant yeast septins apply to other disease-linked mutations in oligomeric proteins.

p53 is a transcription factor and tumor suppressor that is mutated in nearly 50% of human tumors (“IARC p53 Database (p53.iarc.fr)”). WT p53 exists on a precipice of thermostability, where missense substitutions easily push it “over the edge” in terms of its ability to fold, bind DNA and activate transcription (Bullock *et al*. 1997). Tumor-derived p53 mutants vary in conformation and function. In “contact” mutants, DNA binding is specifically abrogated by substitutions in the DNA-binding region without global effects on p53 conformation, whereas in “conformational” mutants the entire DNA-binding domain is misfolded. p53 mutants can also be categorized as dominant or recessive in terms of their inhibitory effect on the function of co-expressed WT p53. Although fungi lack p53, yeast has long been an attractive system for characterizing p53 function using transcriptional reporters whose activity depends on p53 oligomerization, nuclear localization, and DNA binding (Sharma *et al*. 2016; Billant *et al*. 2017). Analysis using a yeast system of 76 mutants representing 54% of over 15,000 total reported missense mutations in the p53 core domain (“IARC p53 Database (p53.iarc.fr)”) found that most were dominant-negative, loss-of-function alleles (Dearth *et al*. 2007). The remaining one-third failed to interfere with WT function and this recessive class was enriched for TS mutants (Dearth *et al*. 2007). We wondered if, like the recessive TS septin mutants, kinetic delays in folding/oligomerization or persistent chaperone engagement play a role in this class of p53 mutants. Coupled with the absence in yeast of many factors that exert extrinsic regulation on p53 oligomerization and turnover (e.g. MDM2 (Di Ventura *et al*. 2008)), the high conservation of all major classes of cytosolic chaperones allows us to focus on effects of mutations on p53 folding, chaperone engagement, and transcriptional activation function.

In this study we set out to determine how chaperone engagement and kinetic delays in folding contribute to the recessive behavior of a TS p53 mutant. Having previously used genetic approaches to find chaperone roles in recessive behavior for septin mutants, we first use mutant septins to test a bimolecular fluorescence complementation (BiFC) screening method to visualize chaperone–client interactions in living cells. We then combine BiFC chaperone screening with genetic and pharmacological manipulation of chaperones to gain a mechanistic understanding of the contributions of protein misfolding and cytosolic chaperones to the ability of a tumor-derived p53 mutant to dominantly interfere with WT function.

## MATERIALS AND METHODS

### Yeast strains, media, plasmids, and genetic manipulations

Yeast strains are listed in Table 1 and were manipulated using standard techniques (Amberg *et al*. 2005). Oligonucleotides are listed in Table 2 and plasmids along with construction details are listed in Table 3. Yeast cells were cultivated in liquid or solid (2% agar) plates of rich or synthetic (Drop Out Mix Minus various ingredients; United States Biological, Salem, MA) medium containing total 2% sugar (glucose, raffinose, or galactose) or 4% sugar (2% each raffinose and galactose), as needed to maintain plasmid selection and induce expression from the *GAL1/10* promoter. The TH5 strain was propagated on rich or synthetic medium with 100 µg/mL dTMP (TCI America #TCT0845), or on minimal medium containing essential amino acids adenine, histidine, and methionine. Drop out medium containing 5-Fluoro-orotic acid (FOA; United States Biological) was prepared with 50 µg/mL uracil and 0.1% FOA, except for use at 37° where modified to 0.07% FOA at pH 4.6. Radicicol (AdiopGen #AG-CN2-0021) was dissolved in DMSO to 10 mg/mL. The bacterial strain DH5α (Agilent Technologies, Santa Clara, CA) was used to propagate plasmids.

**Table 1.**
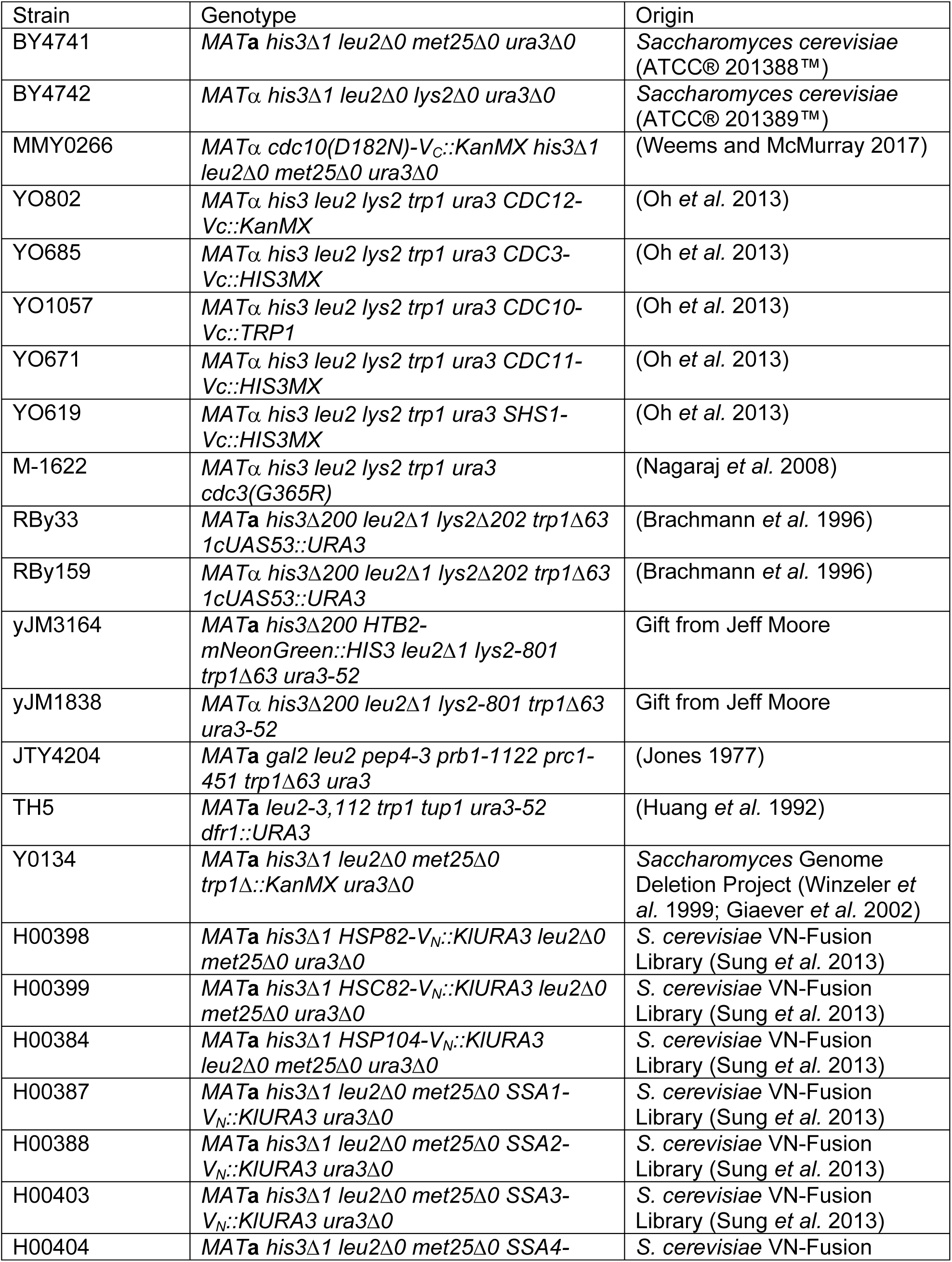
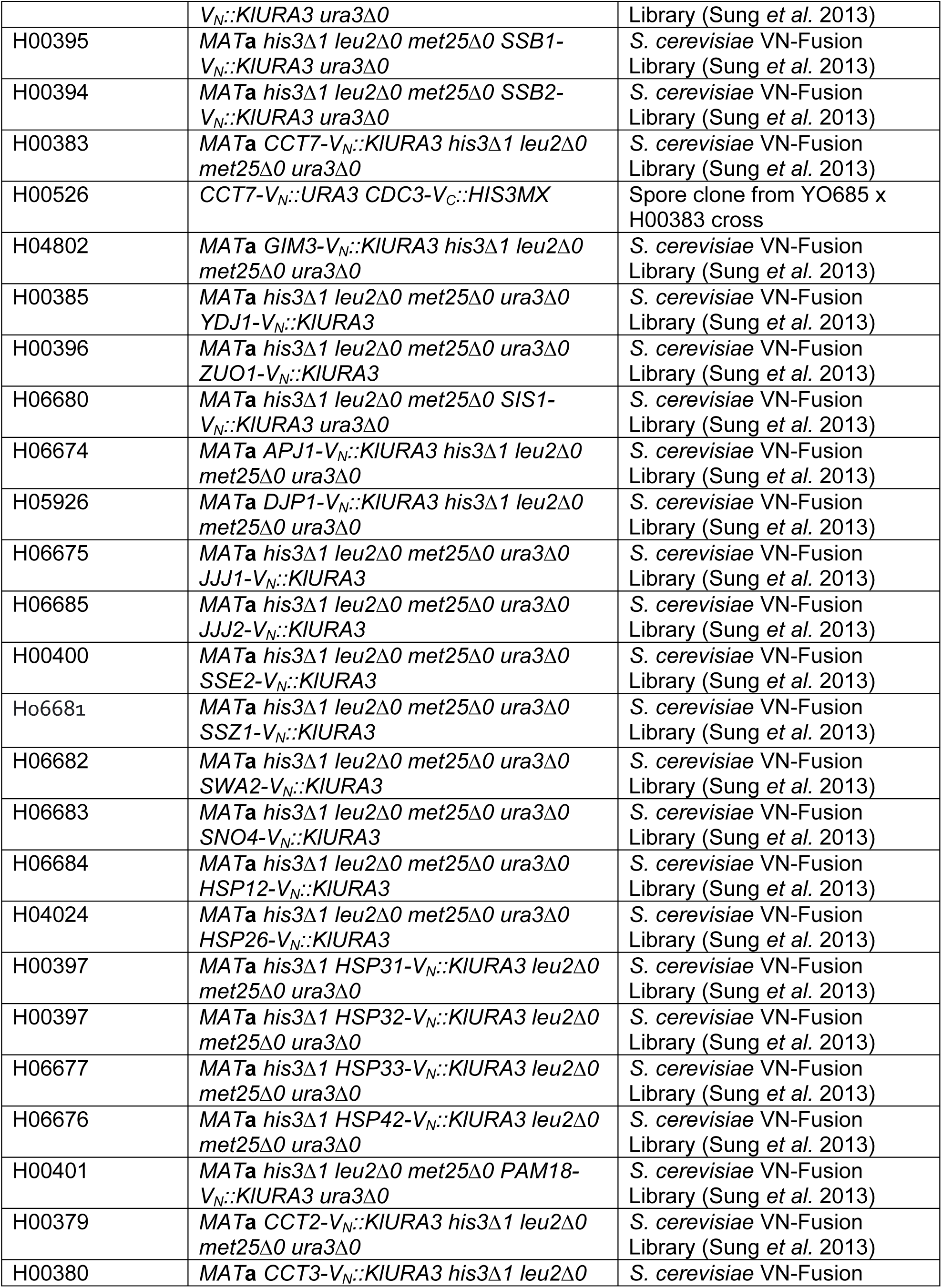
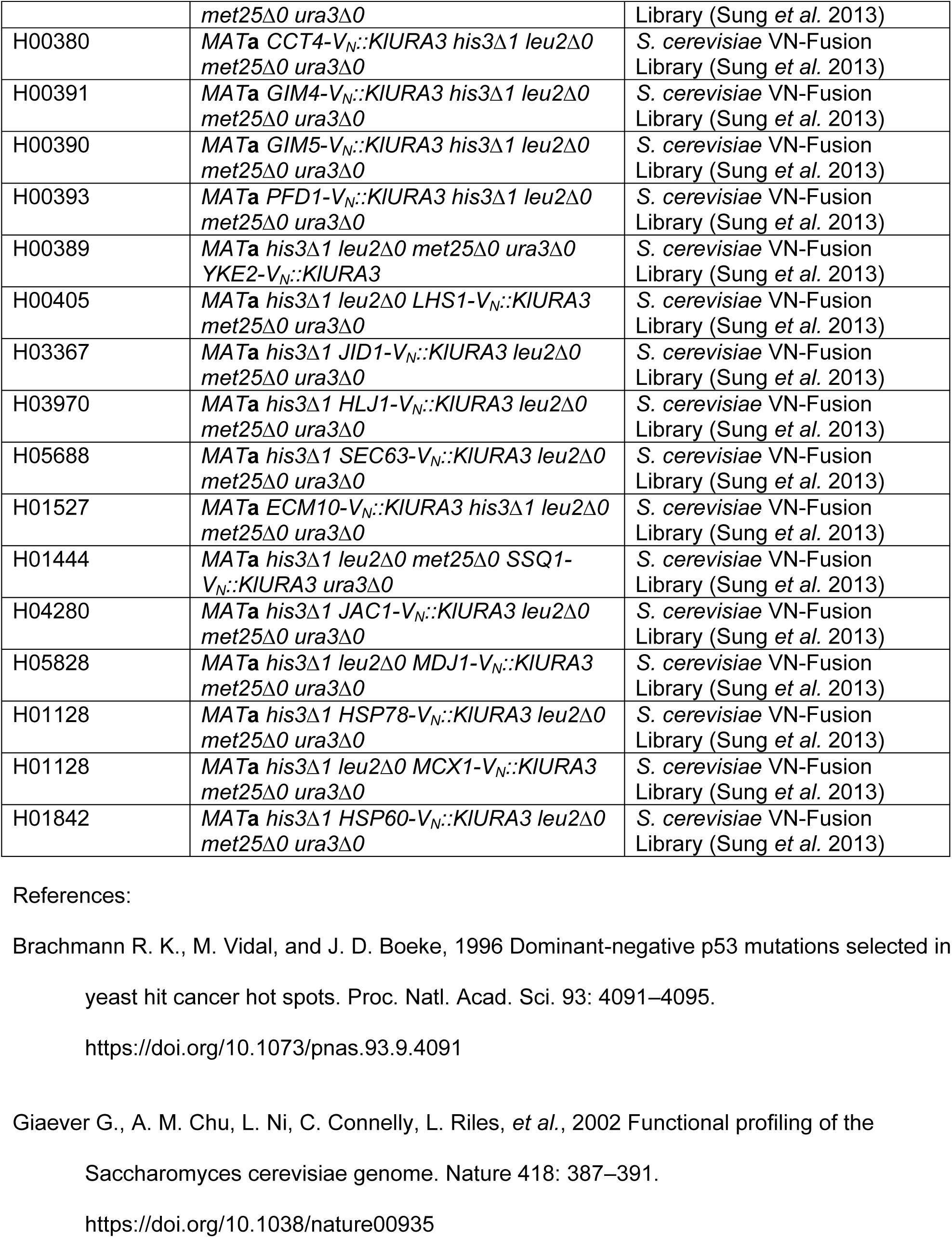
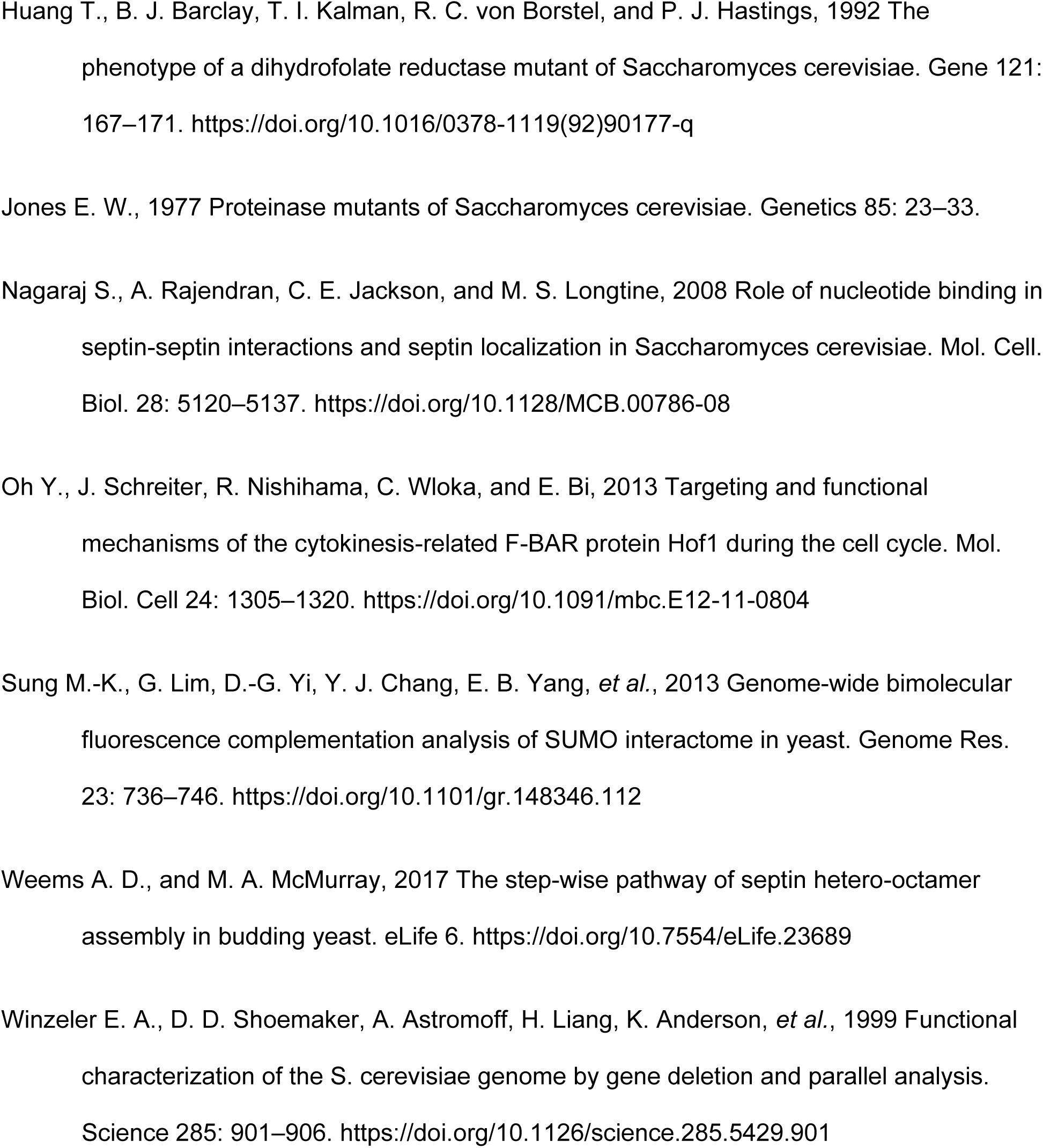
Yeast Strains

**Table 2.**
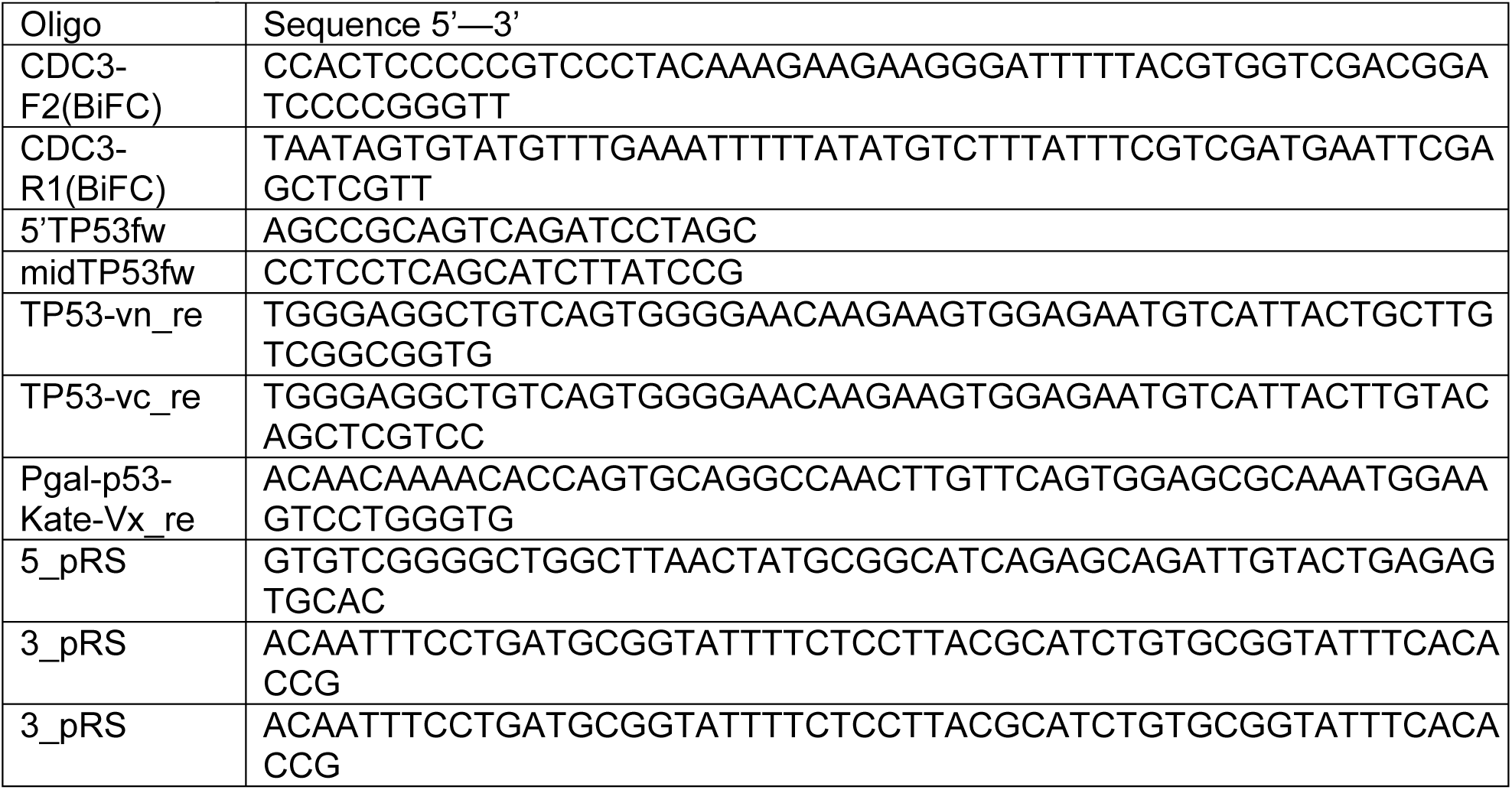
Oligos

**Table 3.**
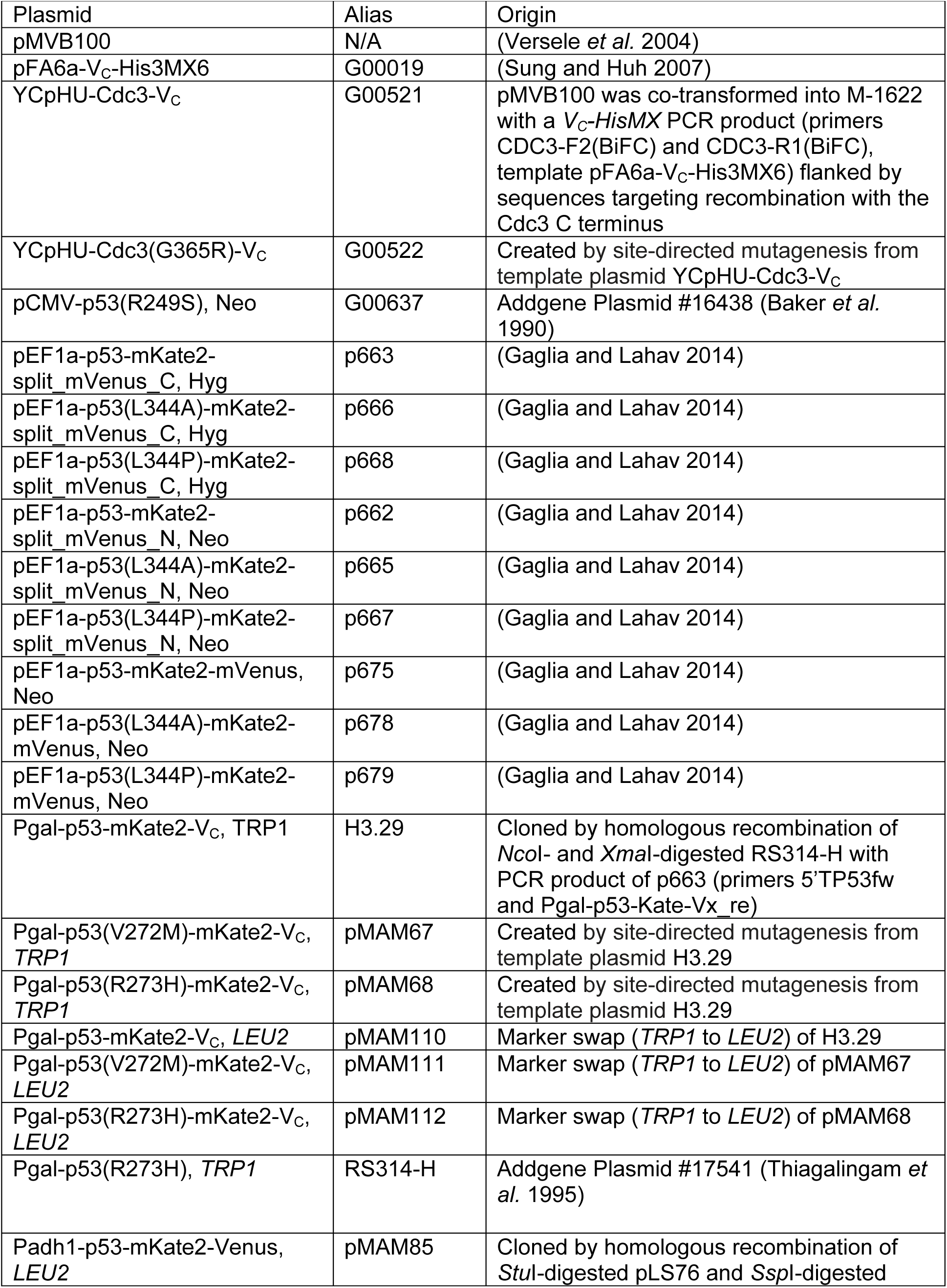
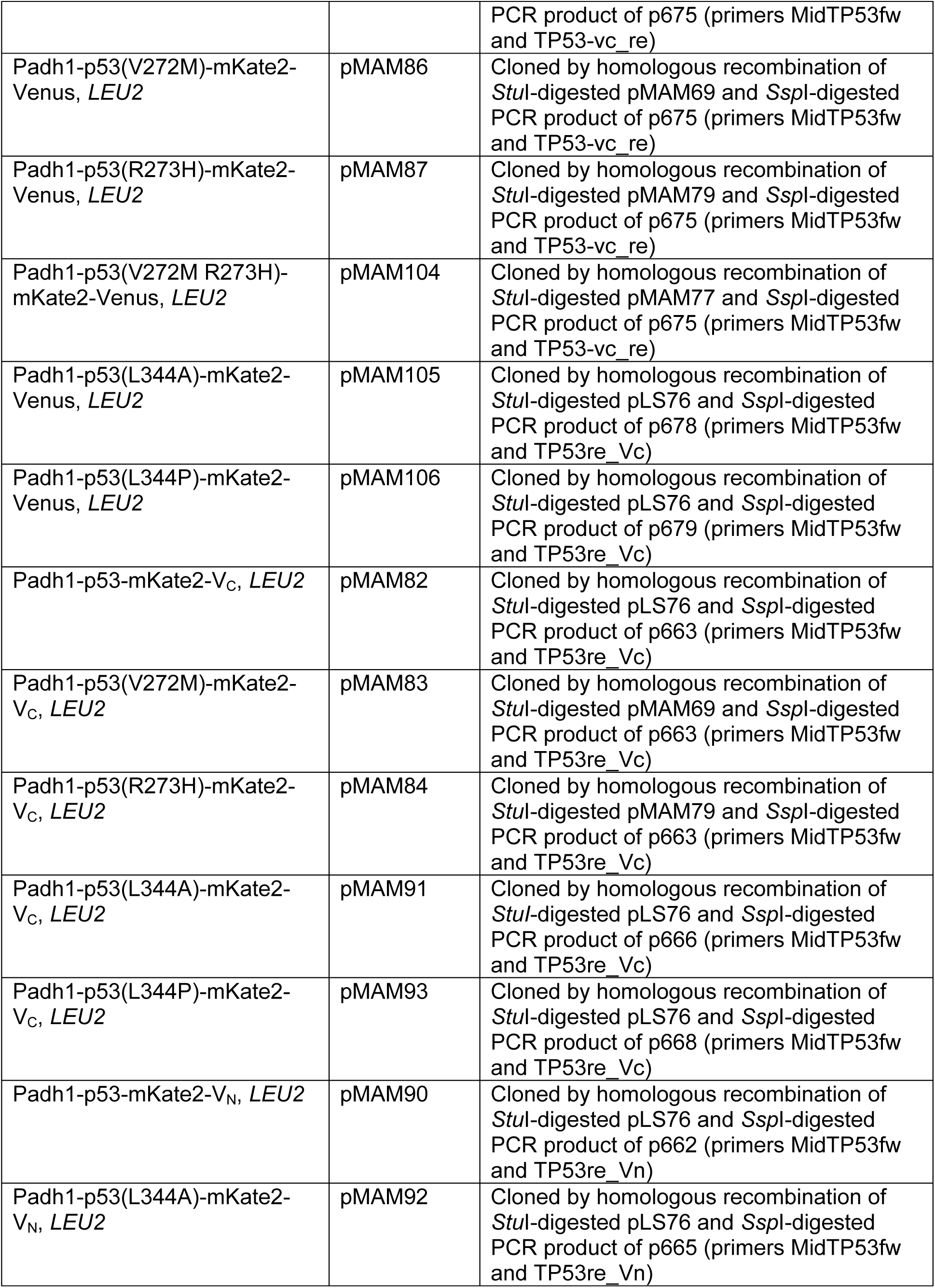
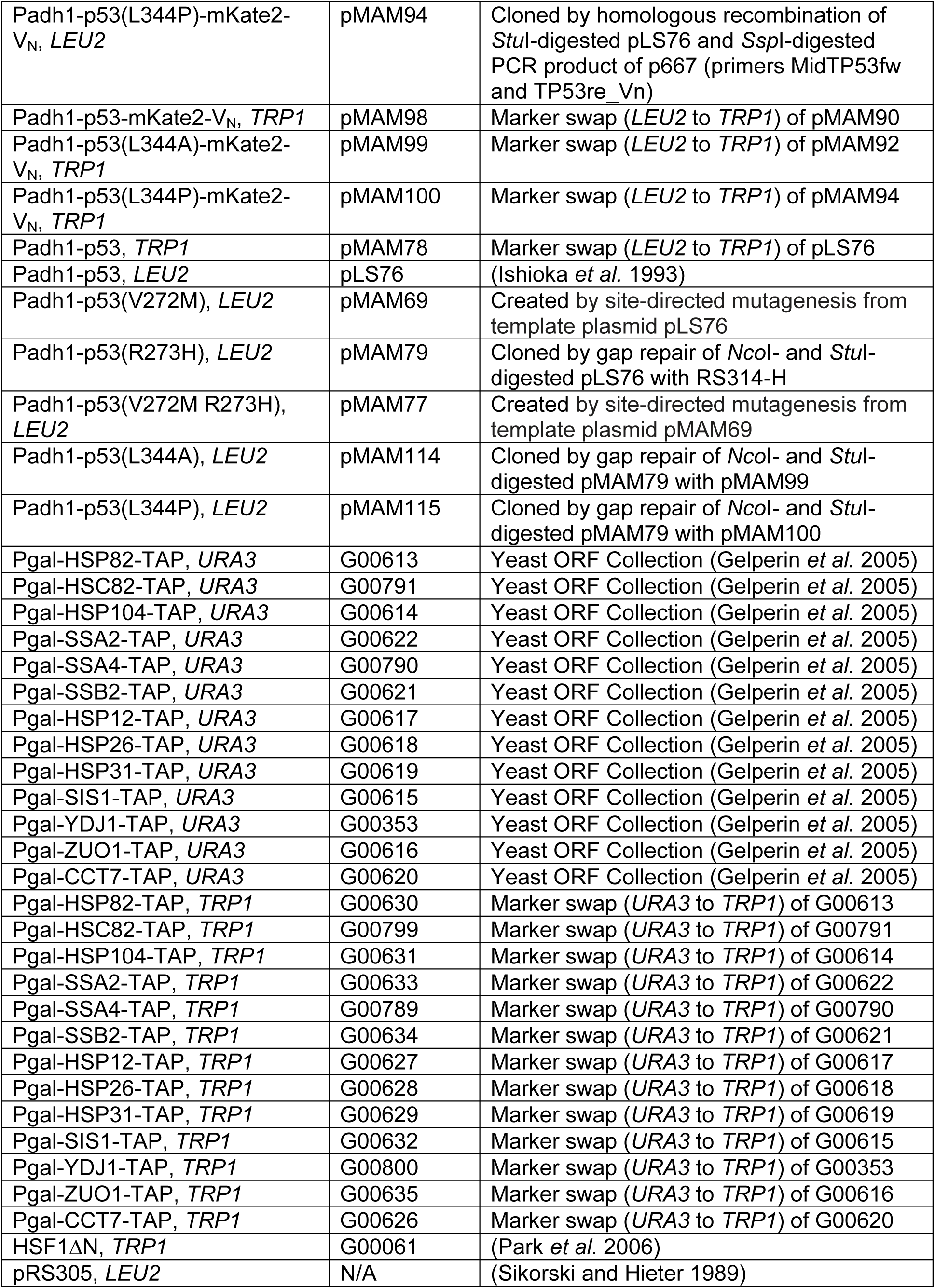
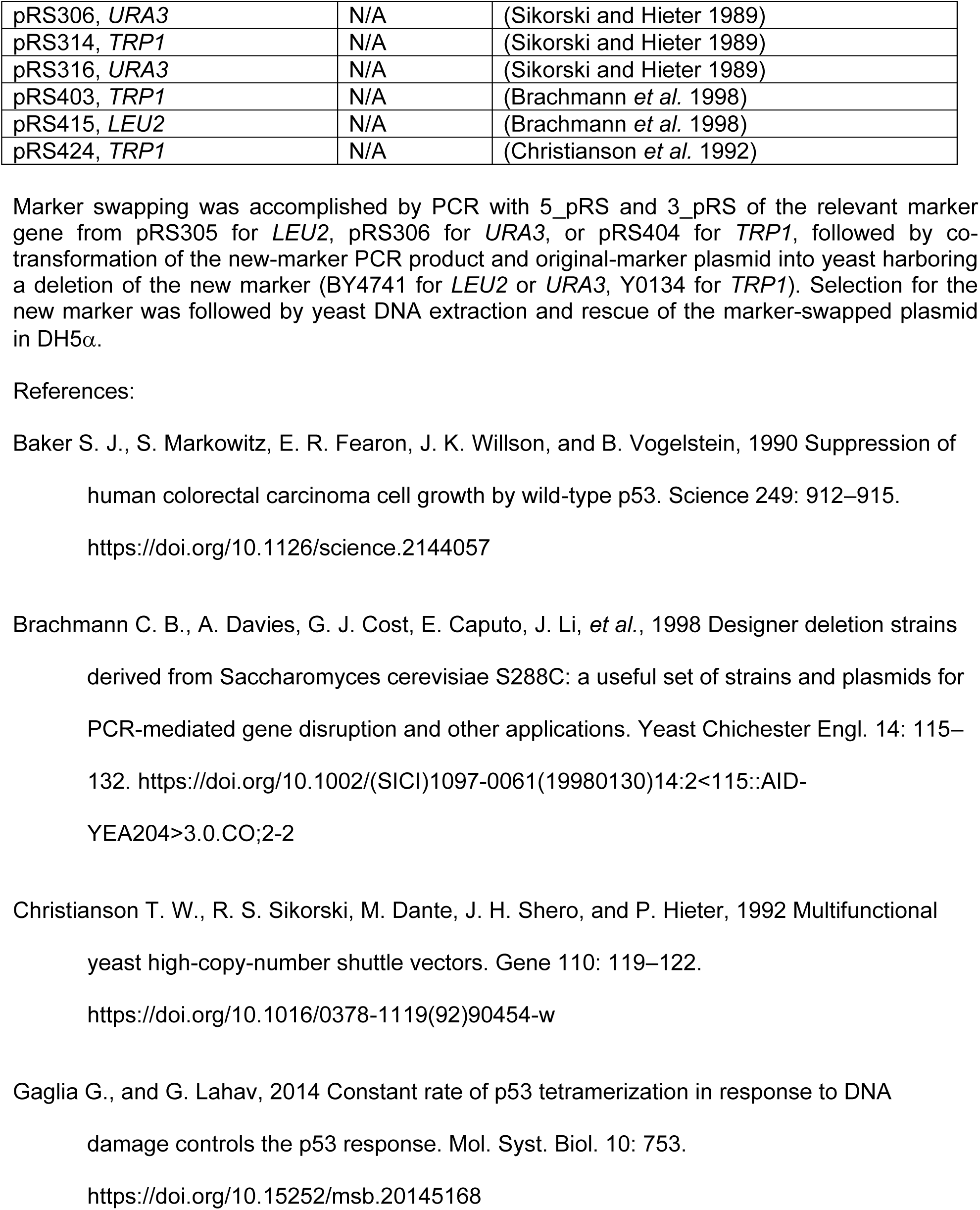
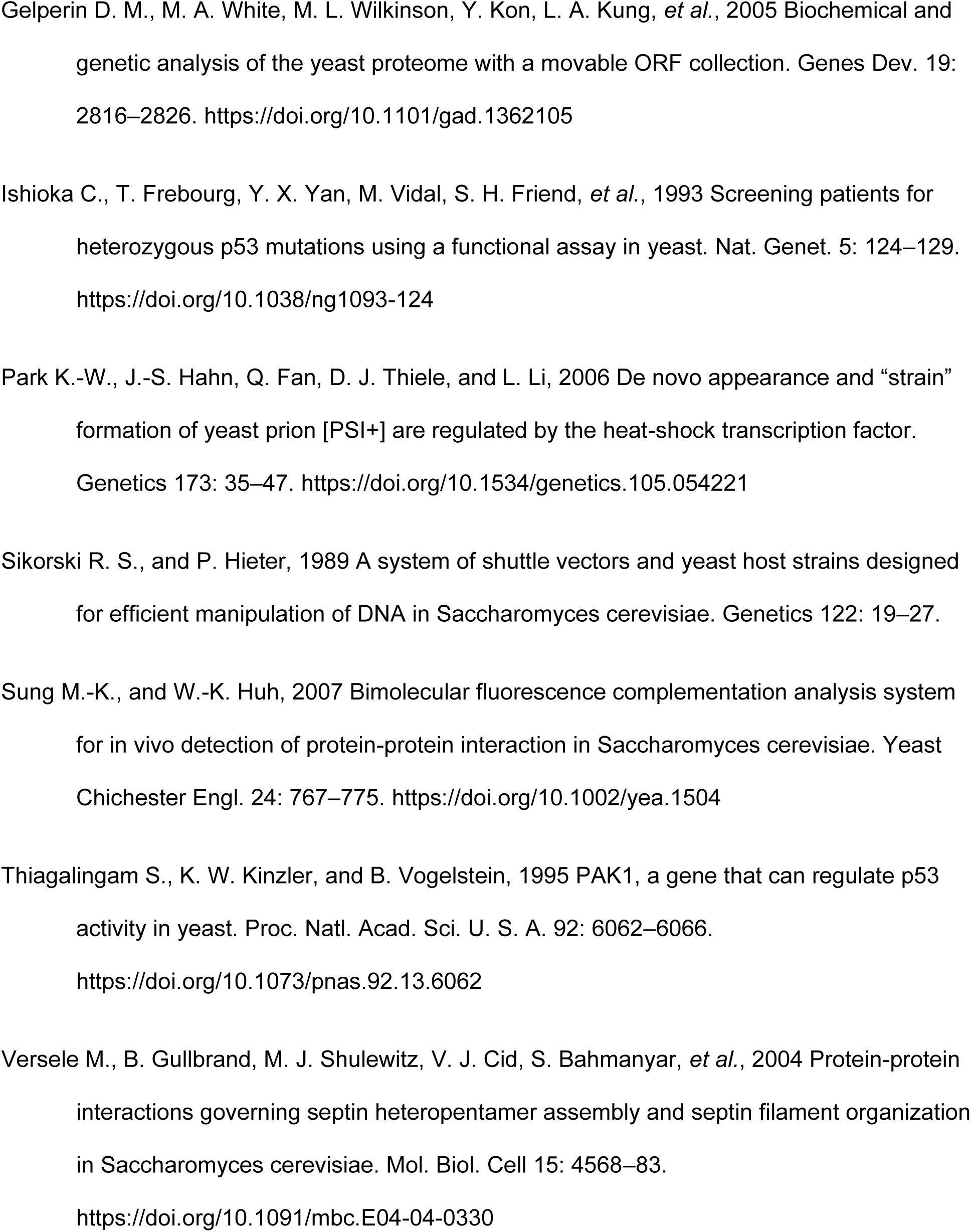
Plasmids

Genomic or plasmid DNA from yeast was isolated by resuspending cells in 250 µL of P1 (50 mM Tris-HCl, pH 8.0, 10 mM EDTA, 100 µg/mL RNAse A), adding 0.5-mm glass beads to displace an additional ∼250 µL of volume, and vortexing for 3 min before addition of 250 µL P2 (200 mM NaOH, 1% SDS) and 350 µL N3 (4.2 M guanidine-HCl, 0.9 M potassium acetate, pH 4.8). Lysates were clarified by centrifugation and applied to Zyppy plasmid DNA Miniprep columns (Zymo Research) and DNA was washed and eluted according to the manufacturer’s instructions. Alternatively, DNA was precipitated from clarified lysates using sodium acetate and ethanol (Green and Sambrook 2016). Yeast transformation was performed using Frozen-EZ Yeast Transformation Kit II (Zymo Research, Irvine, CA). Bacterial transformation was performed using CaCl_2_-prepared competent bacterial cells. PCR was performed with Phusion or Q5 high-fidelity polymerase (New England Biolabs, Ipswich, MA) according to the manufacturer’s instructions. Site-directed mutagenesis was performed by Keyclone Technologies (San Marcos, CA).

### Microscopy

All micrographs were captured with an EVOSfl (Advanced Microscopy Group, Bothell, WA) all-in-one microscope equipped with an Olympus (Tokyo, Japan) 60x oil immersion objective and YFP, Texas Red, and GFP filters. Cells from mid to late log cultures were either taken directly from liquid cultures, applied to glass slides and covered with a coverslip, or concentrated by centrifugation, spotted to agarose pads, and covered with a coverslip. For septin microscopy, LED intensity was 100% and exposure times ranged from 500 to 1500 msec. For p53 microscopy, LED intensity for YFP and Texas Red ranged from 80-100%, and exposure time ranged from 1000 to 2500 msec, together chosen to produce no more than 10% saturated signal. GFP imaging used 20-30% LED intensity and 30-60 msec exposure. For comparison of samples in an experiment all samples were analyzed using the same intensity and exposure time.

### Septin–chaperone BiFC

To measure bud neck BiFC fluorescence (Figure 1C) we used an existing macro (Weems and McMurray 2017) with FIJI software (Schindelin *et al*. 2012). To measure intracellular BiFC signal not focused on the bud neck, we used FIJI to adjust the brightness/contrast on transmitted light images of fields of cells and then used the Threshold function to create a binary image with cell outlines. The “Fill Holes” process was used to fill cell outlines, and then this binary image was used as a mask to measure the integrated signal intensity and areas for the fluorescence image of the same field of cells. To avoid non-cell particles, measurements were restricted to particles with areas between 500 and 2000 square pixels and circularity 0.5-1.0. Occasional cells (<1%) with extremely bright fluorescence were presumed to be dead and were excluded.

**Figure 1.**
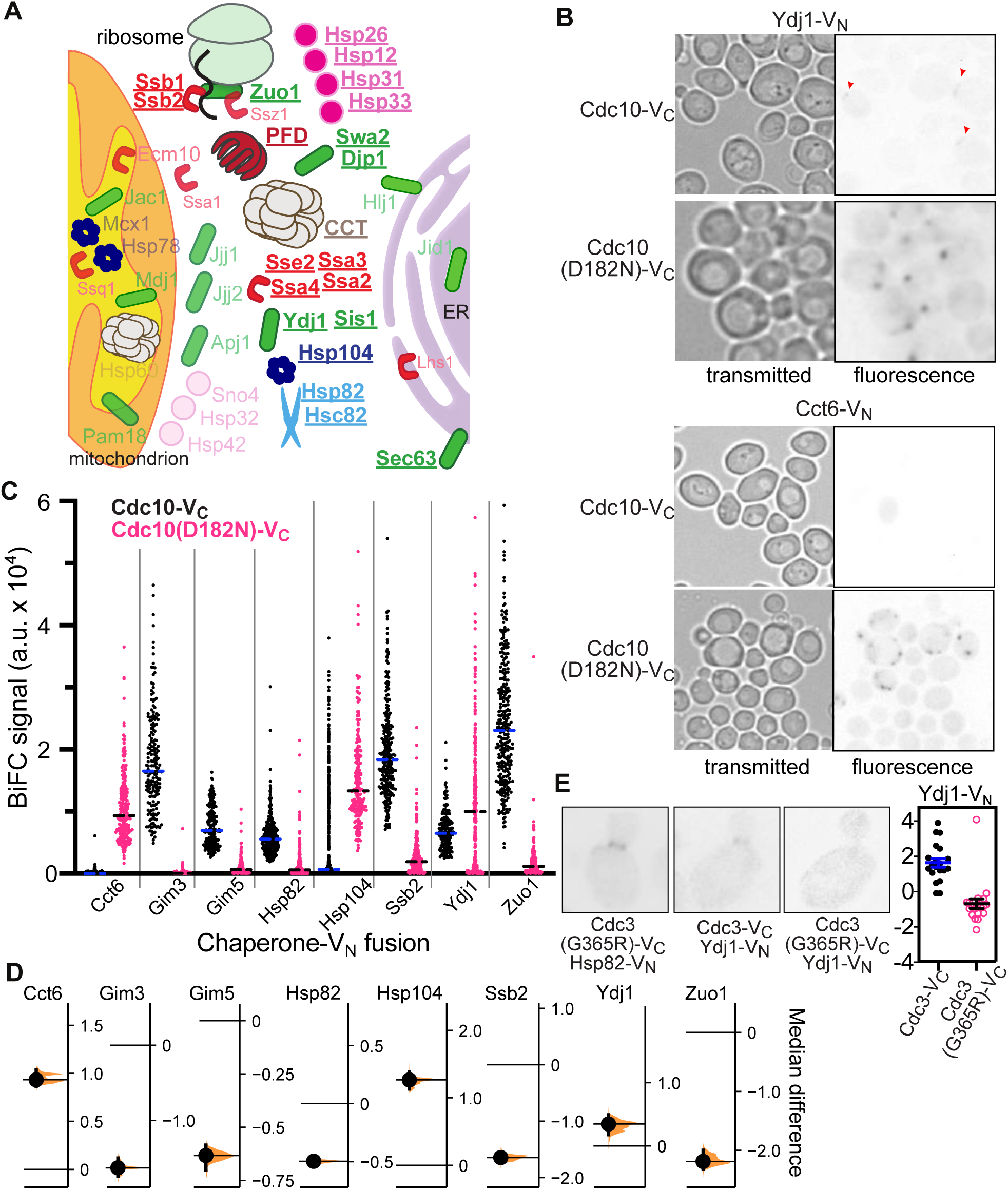
Bimolecular fluorescence complementation reveals septin–chaperone interactions in living cells. (A) Graphical summary of interactions between V_C_-tagged septins and V_N_-tagged cytosolic chaperones in diploid cells co-expressing untagged versions of all proteins. Chaperones that generated discrete fluorescence signal are in bold and underlined. See Table 4 for full dataset. Chaperones are color-coded by family and chaperones from the same family are drawn as the same shape. (B) Micrographs showing discrete fluorescence signals in diploid cells co-expressing Cdc10-V_C_ or the D182N mutant derivative with Ydj1-V_N_ or Cct6-V_N_. The fluorescence images were inverted to facilitate viewing. Red arrowheads point to bud neck signal. Strains were diploids made by mating YO1057 with H00385 or H00382. (C) Cellular BiFC signal away from bud necks was quantified for diploid cells co-expressing Cdc10-V_C_ or Cdc10(D182N)-V_C_ and the indicated V_N_-tagged chaperones. The number of cells analyzed for each genotype ranged from 113 to 460. Strains were diploids made by mating YO1057 or H06530 with H00382, H00392, H00390, H00398, H00384, H00394, H00385, or H00396. (D) For the data in (C), the median difference for comparisons between Cdc10(D182N)-VC and Cdc10-VC are shown in Cumming estimation plots as bootstrap sampling distributions. Each median difference is depicted as a dot. Each 95% confidence interval is indicated by the ends of the vertical error bars. (E) Representative images of diploid cells (strain M-1622 mated with H00398 or H00385) carrying *cdc3(G365R)* at one allele of *CDC3* and WT *CDC3* at the other, a V_N_-tagged allele of the indicated chaperone at one allele of the chaperone gene locus and an untagged allele at the other, and a low-copy plasmid (G00521 or G00522) encoding Cdc3-V_C_ or Cdc3(G365R)-V_C_. Plot at right shows quantification of bud neck fluorescence. 20 cells were analyzed for each genotype.

**Table 4.**
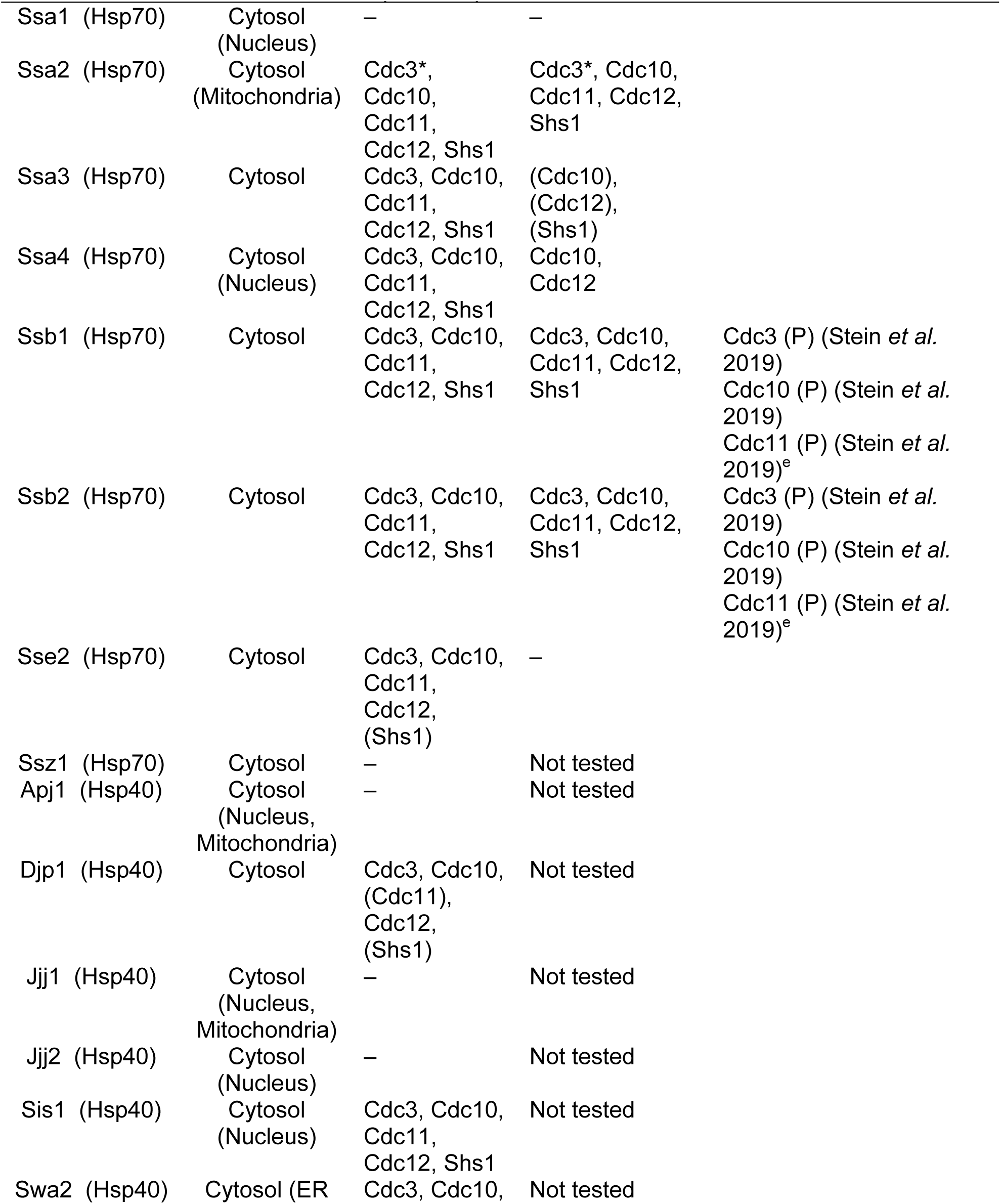
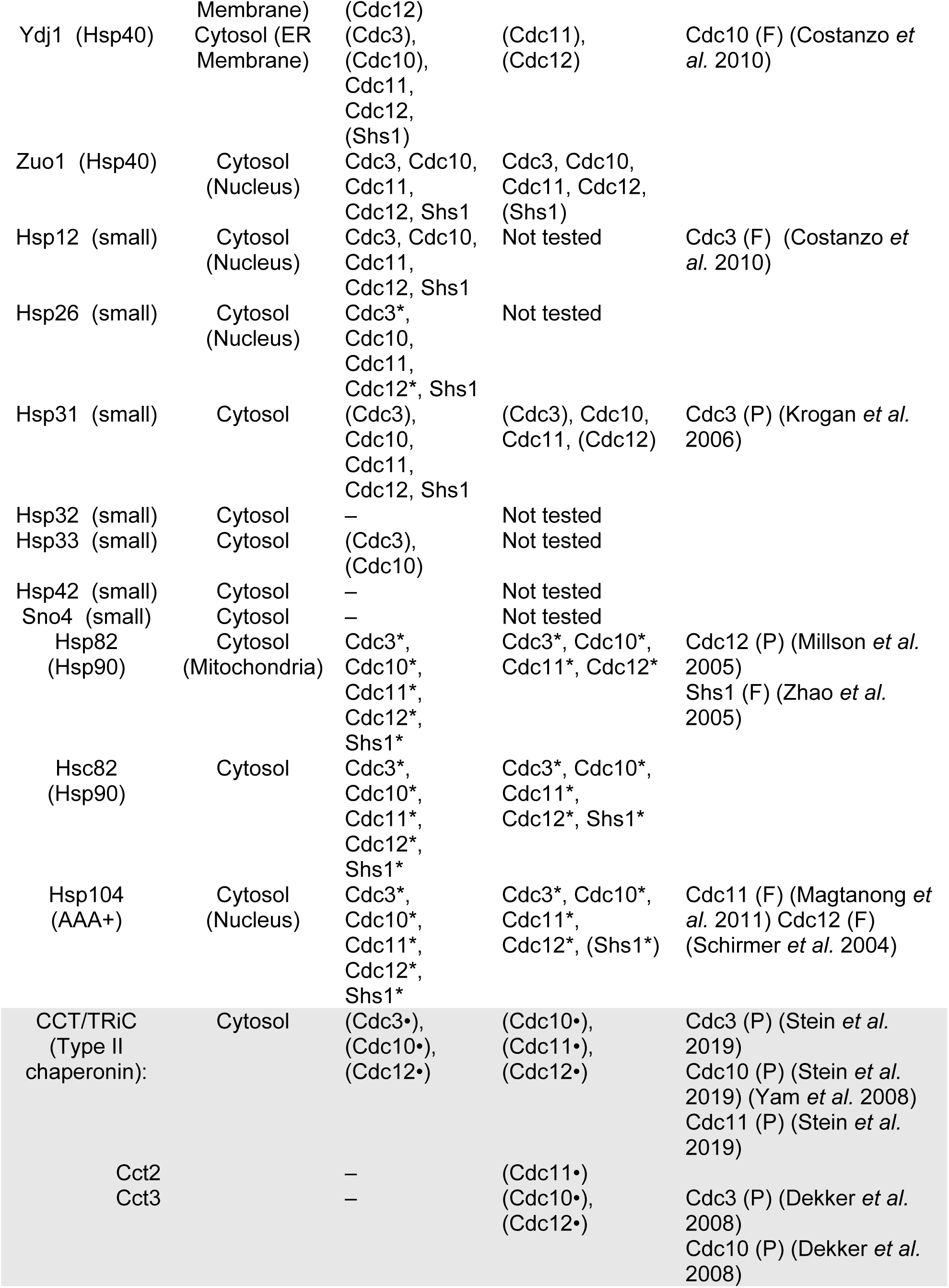
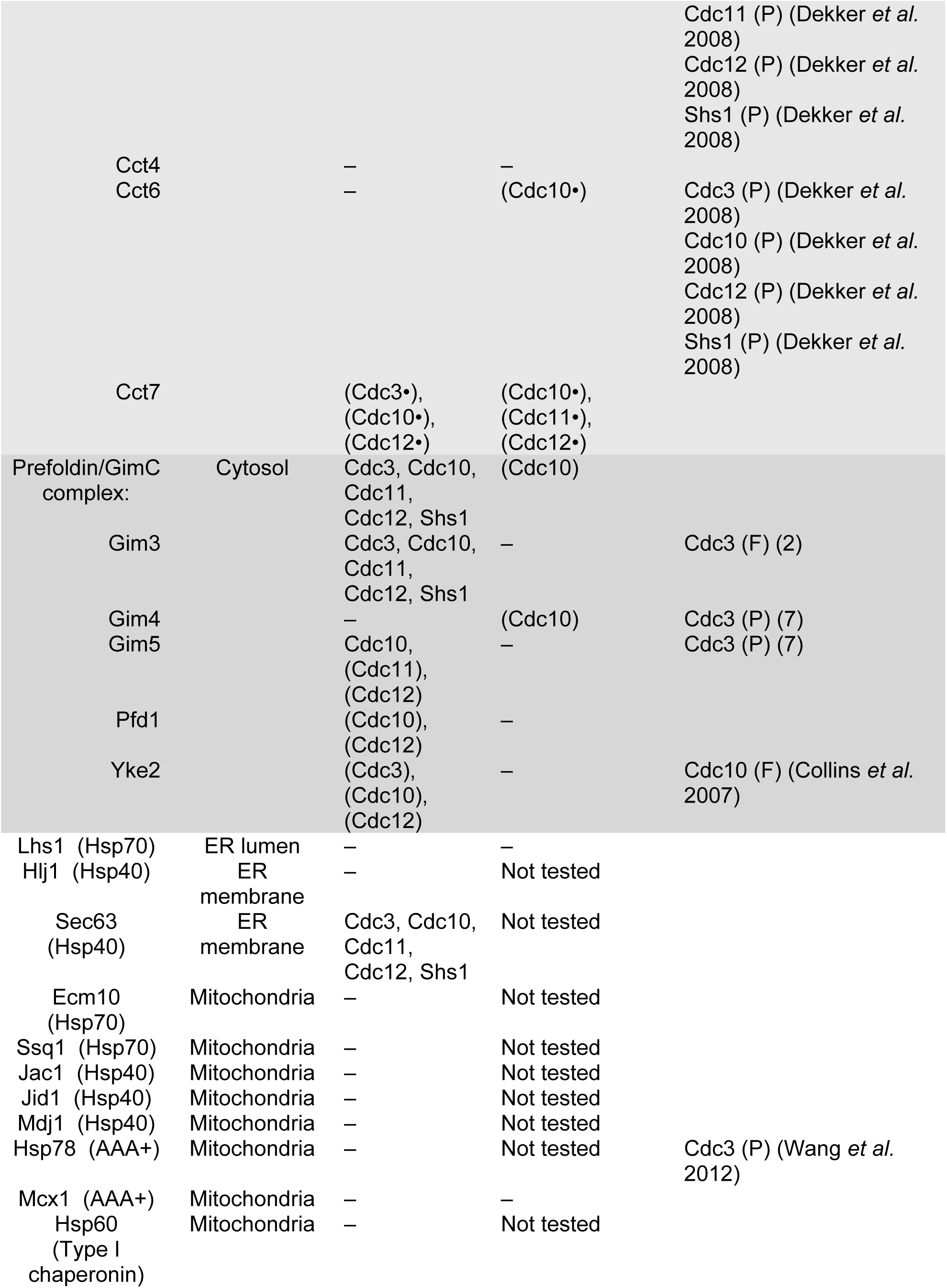
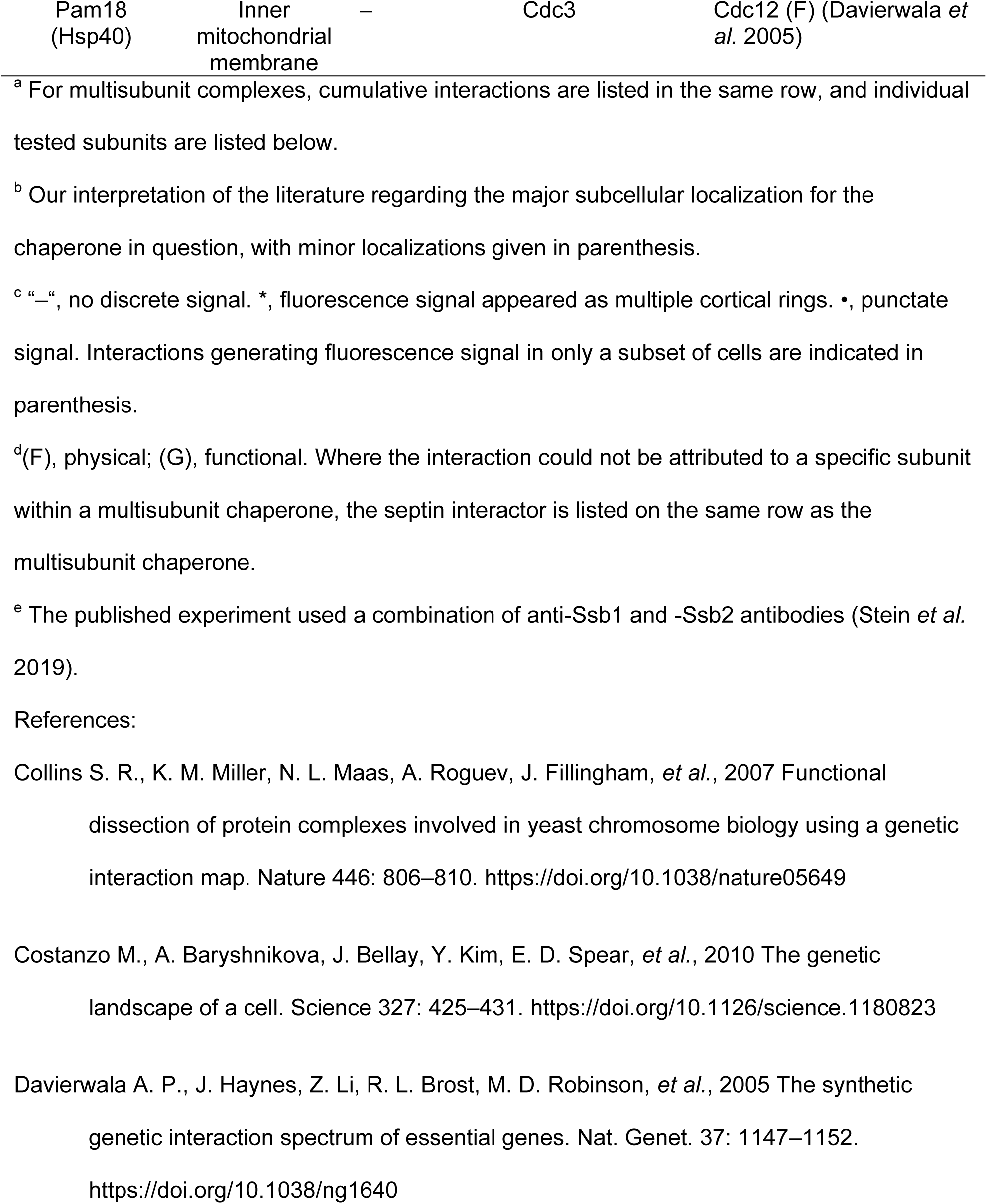
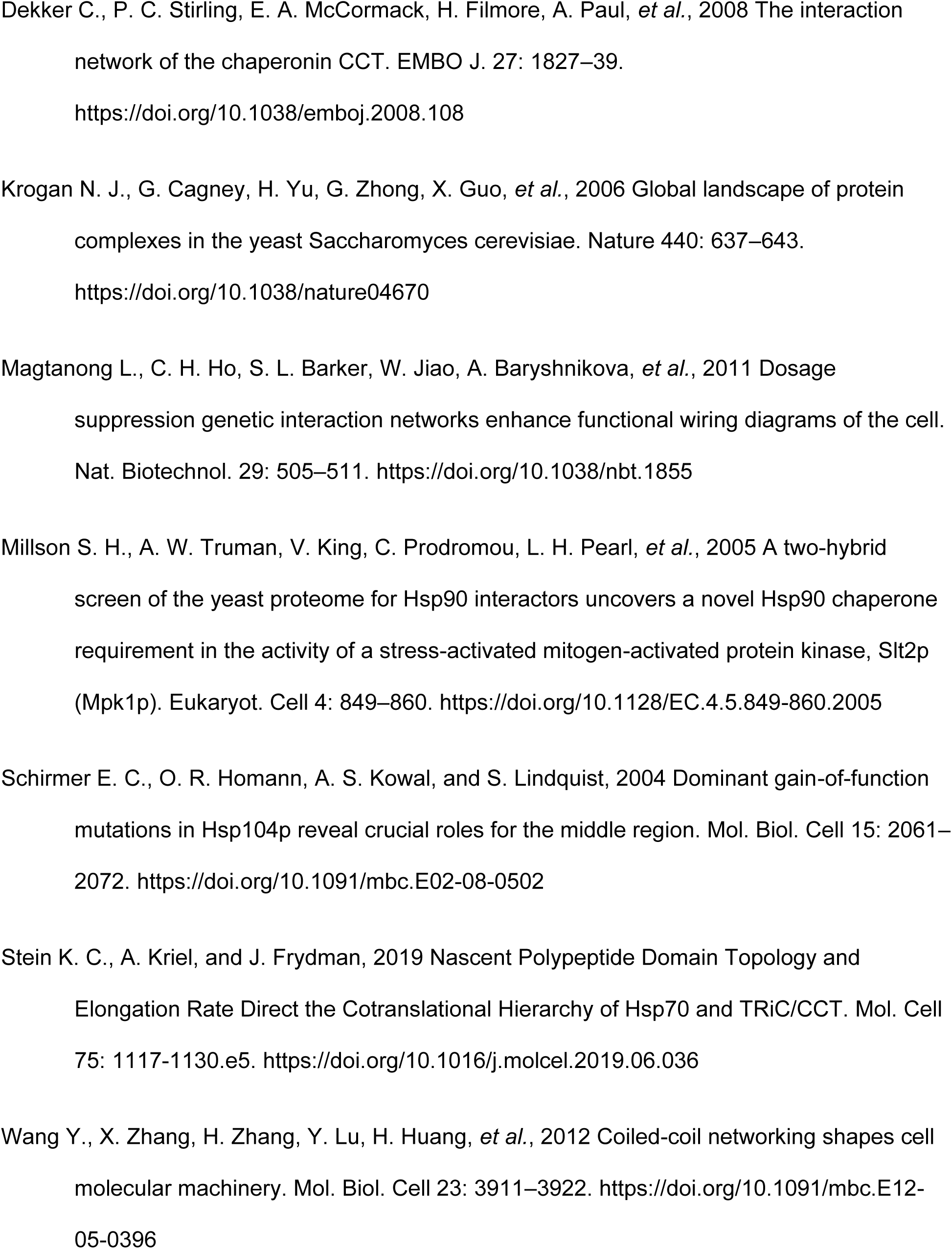
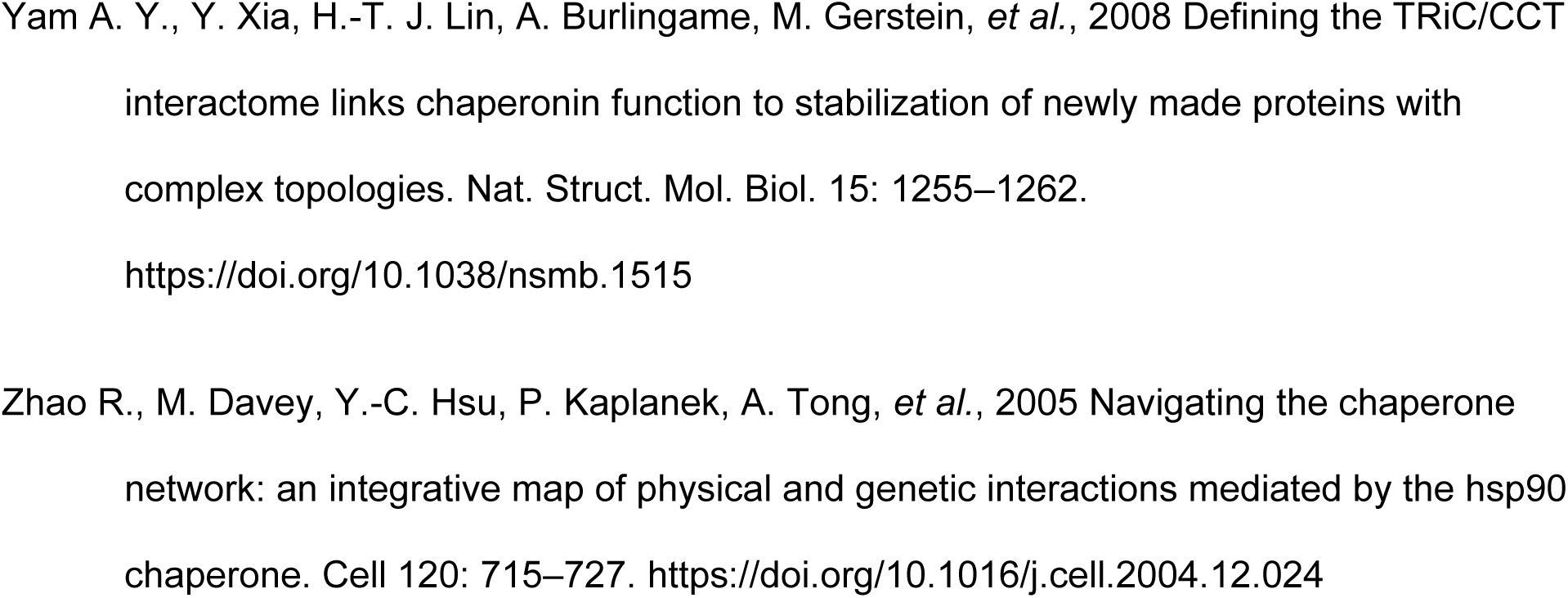
Septin–chaperone interactions assayed by BiFC

### p53–chaperone BiFC

Haploid BY4742 cells carrying relevant plasmids described in figure legends were mated to V_N_-fusion strains encoding a V_N_-tagged chaperone. Diploids were inoculated in liquid synthetic medium with 1.9% raffinose and 0.1% galactose lacking leucine and uracil and imaged after growing to OD_600_∼1 at 30°. For each chaperone the series of WT p53, p53(V272M), and p53(R273H) were imaged in parallel. Micrographs were quantified for Venus and mKate2 signal (n=100 cells for each genotype per chaperone) using FIJI software (Schindelin *et al*. 2012) and a macro that allowed selection and quantification of Venus and mKate2 signals from the same cell location (the cytosol, the nucleus, or both depending on the location of BiFC interactions).

### p53–p53 BiFC

Haploid Y0134 cells carrying relevant plasmids described in figure legends were inoculated in liquid synthetic medium with 2% glucose lacking leucine and tryptophan and imaged after growing to OD_600_∼0.5 at 37°. Micrographs were quantified for nuclear Venus and mKate2 signal (n=100 cells for each genotype) using using FIJI software (Schindelin *et al*. 2012) and a macro that allowed selection and quantification of Venus and mKate2 signals from the same cell location. Cells were selected for quantification based on the presence of Venus signal, as not all cells with mKate2 signal had generated and matured Venus signal, but all cells with Venus signal had corresponding mKate2 signal.

### p53 localization assays

For kinetics of p53 nuclear accumulation, haploid yJM3164 cells carrying relevant plasmids were inoculated from overnight liquid cultures grown at 24° in synthetic medium with 2% raffinose lacking tryptophan into fresh medium with 1.9% raffinose and 0.1% galactose and imaged every 30 minutes for 6 hours at 24°, and finally again at 10 hours. Micrographs were quantified for GFP and mKate2 signal (n=100 cells for each genotype per timepoint) using FIJI software (Schindelin *et al*. 2012) and a macro that allowed selection and quantification of GFP and mKate2 signals from the nucleus. Note that GFP was used here only to verify the location of the nucleus and GFP quantities were not analyzed further.

For steady-state levels of p53 at different temperatures, haploid yJM3164 cells carrying relevant plasmids described in figure legends were inoculated in liquid synthetic medium lacking leucine and imaged after growing to OD_600_∼0.5 at 30° or 37°. Micrographs were quantified for GFP and mKate2 signal (n=100 cells for each genotype per temperature) using FIJI software (Schindelin *et al*. 2012) and a macro that allowed selection and quantification of GFP and mKate2 signals from the nucleus. Note that GFP was used here only to verify the location of the nucleus and GFP quantities were not analyzed further.

For p53 localization with chaperone overexpression, diploid yJM3164 x yJM1838 cells carrying relevant plasmids described in figure legends were inoculated in liquid synthetic medium with 2% galactose and 2% raffinose but lacking leucine and tryptophan and imaged after growing to OD600∼1 at 30°. Micrographs were quantified for GFP and mKate2 signal (n=100 cells per genotype) using FIJI software (Schindelin *et al*. 2012) and a macro that allowed selection and quantification of GFP and mKate2 signals from the nucleus and cytosol. Note that GFP was used here only to verify the location of the nucleus (and by exclusion, the cytosol) and GFP quantities were not analyzed further.

For p53 localization with Hsp90 inhibition, haploid yJM3164 cells carrying relevant plasmids described in figure legends were inoculated in liquid synthetic medium with 2% glucose lacking leucine and containing 5 µg/mL radicicol or equal volume DMSO and imaged after growing to OD_600_∼0.5 at 37°. Micrographs were quantified for nuclear GFP and mKate2 signal (n=100 cells per genotype) using FIJI software (Schindelin *et al*. 2012) and a macro that allowed selection and quantification of GFP and mKate2 signals from the nucleus and cytosol. Note that GFP was used here only to verify the location of the nucleus (and by exclusion, the cytosol) and GFP quantities were not analyzed further.

### Co-IP and immunoblotting

25-mL yeast cultures in liquid synthetic medium lacking leucine and tryptophan were grown to OD_600_∼1 at 30° and pelleted. Pellets were washed with Lysis Buffer (20 mM Tris-HCl pH 7.4, 150 mM NaCl, 1 mM EDTA, 0.5% Triton X-100, 1 mM DTT) plus protease inhibitors (Complete EDTA-free; 11 873 580 001, Roche) and frozen in liquid nitrogen. Pellets were resuspended in 300 µL ice-cold Lysis Buffer plus protease inhibitors and lysed using 100 µL glass beads and 8 cycles of 30-second on/off vortexing at 4°. Lysates were clarified by centrifugation into new tubes. Clarified lysates were diluted 1:2.5 in Binding Buffer (10 mM Tris-HCl pH 7.4, 150 mM NaCl, 0.5 mM EDTA, 1 mM DTT) for a total volume of 700 µL “Input”. 25 µl Chromotek GFP-nAb Magnetic Agarose beads (Chromotek, Germany) were washed three times with Binding Buffer and combined with 700 µL diluted lysate. Beads were incubated while tumbling end-over-end for 1 hour at 4°. “Unbound” fraction was removed and beads washed once with Binding Buffer and twice with Wash Buffer (10 mM Tris-HCl pH 7.4, 500 mM NaCl, 0.5 mM EDTA, 1 mM DTT). Beads were resuspended and gently pipetted in 50 µL 2x SDS–PAGE sample buffer plus 5% β-mercaptoethanol and heated at 95° for 5 min to generate the “Bound” fraction. Following SDS-PAGE, proteins were electrophoretically transferred to PVDF and exposed to mouse anti-p53 (RRID:AB_10986581) and rabbit anti-Pgk1 primary antibodies. Infrared-labeled anti-mouse (680 nm, RRID:AB_10854088) and anti-rabbit (800 nm, RRID:AB_10697505) secondary antibodies were applied before scanning on a Licor Odyssey device. Input represents 7% of the amount of total protein used for the amount of immunoprecipitated material loaded on the gel.

For all other immunoblots, cultures were grown to mid-log phase and pellets were washed with 10% trichloroacetic acid (TCA) and frozen in liquid nitrogen, then placed at -70°. Cell pellets were thawed in 150 μL of 1.85 M NaOH and 7.4% β-mercaptoethanol. 150 μL of 50% TCA was added before incubation on ice for 10 min. Precipitated proteins were pelleted at 4° at maximum speed for 10 min, washed twice with cold acetone (1 mL each time), and resuspended in 80 μL of 0.1 M Tris base, 5% SDS. After addition of SDS–PAGE sample buffer, the samples were boiled for 5 min before resolution by SDS–PAGE. Membrane processing was the same as described above and also utilized, where indicated, mouse anti-6xHis antibodies (UBPBio #Y1011). Ponceau solution (0.5% w/v, 1% acetic acid), where indicated, was applied for 30 minutes, then washed with water and the membrane allowed to dry before imaging.

### Choice of sample sizes and statistical analysis

Sample sizes (number of cells) for fluorescence microscopy were chosen based on our ability to detect changes in subcellular localization in our previous studies (Johnson *et al*. 2015, 2020; Weems and McMurray 2017). Analysis of effect sizes and generation of effect size plots were performed using DABEST (data analysis with bootstrap-coupled estimation) (Ho *et al*. 2019) via the online server at http://www.estimationstats.com.

### Data availability

Strains and plasmids are available upon request. The authors affirm that all data necessary for confirming the conclusions of the article are present within the article, figures, and tables.

## RESULTS

### Bimolecular fluorescence complementation (BiFC) as an approach to trap otherwise transient chaperone–client interactions

In principle, the chaperone–septin interactions we found using biochemical approaches (Johnson *et al*. 2015) could have occurred following cell lysis. To visualize chaperone–septin interactions in intact, living cells, we used a BiFC approach, also called split-YFP. Reconstitution of Venus, a yellow fluorescent protein (YFP) derivative, occurs when two non-fluorescent Venus fragments, V_N_ and V_C_, are brought in close proximity. Fusion of V_N_ and V_C_ in parallel to two proteins that directly interact *in vivo* is typically sufficient to drive Venus reconstitution. Although the interactions driving its folding and chromophore maturation are non-covalent, Venus reconstitution is effectively irreversible *in vivo*. Consequently, BiFC can be used to generate a stable fluorescent signal from otherwise transient interactions in which the untagged proteins would normally dissociate (Hu *et al*. 2002). Consequently, BiFC can be used to generate a stable fluorescence signal from otherwise transient interactions. (Chromophore maturation requires considerable additional time after Venus folding, up to 50 minutes (Kerppola 2008), hence signal intensity over time also depends on the half-life of the “crosslinked” fusion proteins.) Since chaperone–client interactions are transient by definition, BiFC is ideally suited for detecting such interactions *in vivo*.

Venus fragments are misfolded proteins that could act as “chaperone bait” independent of the protein to which they are fused. Indeed, multiple V_C_-tagged chaperones (Hsp12, Hsc82, Hsp82, Hsp104, Ssa1, Ssb1, Ssb2, Sec63, Sis1, and Zuo1) are known to generate BiFC signals when co-expressed with “free” V_N_ (Kim *et al*. 2019). We reasoned that the smaller size of V_C_ (∼120 residues, ∼13.5 kDa) compared to V_N_ (∼190 residues, ∼21 kDa) might make it less prone to non-specific chaperone interactions. We therefore mated haploid yeast cells expressing V_C_-tagged septins with haploid cells expressing V_N_-tagged chaperones and after culturing the resulting diploid cells at room temperature (∼22°) we examined them by epifluorescence microscopy. As shown in Figure 1 and summarized in Figure 1A and Table 4, we observed discrete fluorescence signals for multiple cytosolic chaperones. In most cases these signals were found in either of two general subcellular locations. While some diffuse BiFC signal was also faintly visible (see below), we focused first on the much more obvious discrete signals.

The first location of BiFC signal, observed with numerous chaperones, resembled the typical localization observed for native septins: rings at the cortex (Figure 1B, Figure S1A, Table 4). Cortical rings were seen either exclusively at the bud neck, where septin rings are normally found, or faintly at bud necks and more intensely at locations reminiscent of bud-site-selection landmark proteins that persist at previous sites of budding (Figure S1A). In unstressed cells, most fluorescently-tagged cytosolic chaperones are found distributed diffusely throughout the cytosol (Huh *et al*. 2003). We think it is most likely that for the chaperones generating cortical ring fluorescence, BiFC events arose from transient septin–chaperone interactions in the cytosol. The tagged chaperone was then forced to follow the tagged septin on its subsequent cortical journeys. We interpret BiFC signal at bud necks as representing “crosslinked” septin-V_C_- chaperone-V_N_ complexes in which the septin incorporated into septin hetero-oligomers and then into septin filaments, with the tagged chaperone becoming tethered to the septin ring. Due to the delay between initial interaction and fluorophore maturation, by microscopy we see only “old” septin–chaperone “crosslinked” complexes that may have already been through multiple rounds of filament polymerization and depolymerization. The appearance of bright cortical rings at sites of prior budding for some chaperones may indicate that those specific septin-chaperone BiFC associations perturbed the disassembly of “old” septin rings that normally follows each cell division, such that new rings are dimmer because they preferentially incorporate “new” or untagged septins.

The second type of discrete BiFC signal was unique to interactions between WT septins and subunits of the cytosolic chaperonin called CCT (chaperonin containing tailless complex polypeptide 1). Co-expression of Cct7-V_N_ with Cdc3-V_C_ Cdc10-V_C_, or Cdc12-V_C_ generated discrete foci in a subpopulation of cells (Figure S1A and Table 4). We engineered haploid strains carrying Cct7-V_N_ and Cdc3-V_C_ and confirmed that the punctate BiFC signal was not unique to diploid cells (Figure S1A). That these haploids were viable and of normal morphology indicates that the septin–Cct7 BiFC interaction did not perturb essential CCT functions, and that only some of the Cdc3-V_C_ molecules underwent BiFC events, otherwise no Cdc3-V_C_ would have been available to perform septin functions at the bud neck. In stressed cells, other chaperones (e.g., Hsp104) localize to punctate structures together with misfolded proteins, such as the IPOD (insoluble protein deposit) and/or JUNQ (juxtanuclear quality control) (Kaganovich *et al*. 2008), but the Cct7–septin BiFC foci did not co-localize with Act1(E364K)-mCherry, a misfolded form of actin that marks these compartments (Figure S1B). CCT subunit C termini project into the center of the chamber (Llorca *et al*. 1999), hence Cct7-V_N_– septin-V_C_ interactions likely occur when septins are inside, as opposed to interacting with the chamber surface. We cannot exclude the alternative possibility that Cct7-V_N_ interacts with septins prior to assembling into a 16-mer. Although each individual CCT subunit is relatively small, the cage-shaped holoenzyme is large (∼1 MDa). We interpret the appearance of foci as evidence that following BiFC association the tagged septin remains tethered to the inside of the chamber, and is unavailable for incorporation into septin-based structures. CCT assemblies with septins tethered via BiFC may be unable to fold other proteins, but why they would accumulate together in a single intracellular location is a mystery. The apparent ability of Cdc3-V_C_ to frequently “escape” Cct7-V_N_ interaction could mean that some Cdc3 never interacts with CCT, that some Cdc3–CCT interactions are too transient for BiFC, and/or that Cdc3–CCT interactions do not always involve the Cct7 subunit.

Individual CCT subunits make client-specific contacts. For example, actin directly contacts only five CCT subunits (Balchin *et al*. 2018). When CCT is dissociated into individual subunits, Shs1 co-purifies with Cct6 but not with Cct3 (Dekker *et al*. 2008), suggestive of some specificity in CCT–septin interactions. Indeed, Cct7-V_N_ interacted with Cdc3-V_C_, Cdc10-V_C_ and Cdc12-V_C_, but did not exhibit a BiFC signal with Shs1-V_C_ or its paralog Cdc11-V_C_ (Table 4). We did not detect punctate septin BiFC signal with V_N_-tagged Cct2, Cct3, Cct4, or Cct6 in cells cultured at room temperature; Cct1/Tcp1, Cct5, and Cct8 are not in the V_N_ tag collection and thus were not tested. Another multisubunit chaperone complex, prefoldin, also displayed subunit-specific BiFC interactions with septins (Table 4).

Temperature affects both protein folding and the expression of most chaperones. To ask if our BiFC system was able to capture such changes, we repeated our analysis in cells cultured at 37°. The spectrum of chaperone interactions at this temperature was mostly unchanged, with a few notable exceptions. At 37° Cct2, Cct3, and Cct6 showed detectable signal, as discrete puncta similar to what we saw for Cct7 at room temperature (Table 4 and data not shown). Other septin–chaperone BiFC signals decreased at 37°. Septin interactions with prefoldin subunits almost completely disappeared at 37° (Table 4). Interactions between some septins (e.g., Cdc10) and the Hsp40 Ydj1 were no longer detectable, and interactions with several cytosolic Hsp70- family chaperones were notably reduced (Table 4). Since many other chaperone interactions, including Hsp70s, were unchanged, an intrinsic reduction in the efficiency of Venus BiFC at 37° cannot explain the loss of specific chaperone signals. Instead, we attribute these changes to effects of temperature on septin folding – exposing different chaperone-interacting sequences – and/or chaperone expression levels.

### Unique interactions between cytosolic chaperones and mutant septins that are kinetically delayed in their folding

Having established that WT septins associate with cytosolic chaperones, we wished to test a prediction from our KADO observations: that a slow-folding mutant septin would display distinct chaperone interactions relative to the WT allele. We previously showed that the D182N substitution in the GTP-binding pocket of septin Cdc10 renders the mutant protein unable to compete with WT Cdc10 for incorporation into septin complexes, and increases association with the chaperones Hsp104 and Ydj1, according to co-fractionation and co-precipitation using cellular lysates (Johnson *et al*. 2015). To compare chaperone interactions using BiFC, we introduced the D182N mutation into the Cdc10-V_C_ strain and mated with strains expressing V_N_-tagged chaperones.

In no case did we observe fluorescence in septin rings, which presumably reflects KADO-mediated exclusion of the mutant septin from septin complexes, and instead for some chaperones intracellular fluorescence was observed in other locations, including diffusely throughout the cells or in puncta (Figure 1B and data not shown). To compare chaperone interactions between WT Cdc10 and Cdc10(D182N), we measured intracellular fluorescence using cell outlines, which mostly ignored cortical septin ring signals in WT cells (Figure S1C). Several differences were obvious. Whereas WT Cdc10-V_C_ generated robust signal with Gim3-V_N_, almost no fluorescence was detected with Cdc10(D182N)-V_C_ (Figure 1C,D). Due to a lack of discrete fluorescence signal, we had not previously called Gim5, another prefoldin subunit, as a septin interactor (Table 4), but quantification revealed diffuse intracellular fluorescence with Cdc10-V_C_ that decreased drastically for Cdc10(D182N)-V_C_ (Figure 1C,D). Conversely, little to no BiFC signal was found in cells co-expressing Cdc10-V_C_ and Cct6-V_N_ (Figure 1B,C,D), in agreement with our qualitative calls based on discrete signals (Table 4), but Cdc10(D182N)-V_C_ generated much higher signals, often visible as multiple puncta per cell (Figure 1B,C,D). BiFC signal decreased drastically for the cytosolic Hsp90 isoform Hsp82, but not the more highly expressed isoform Hsc82 (Figure S1D). BiFC signal also decreased for Ssb2 and Zuo1 (Figure 1C,D), which comprise an Hsp70–Hsp40 pair within the ribosome-associated complex (RAC) of chaperones that engage nascent polypeptides and promote co-translational folding (Gautschi *et al*. 2002; Preissler and Deuerling 2012; Zhang *et al*. 2017). For the Hsp40 chaperone Ydj1 the change in BiFC signal was more complex. Two populations of cells, differing in the intensity of BiFC fluorescence, were apparent in cultures of each genotype, but the difference was exaggerated by the D182N mutation in Cdc10, with about half of the cells exhibiting almost no signal and the other half having signal higher than the median signal for WT Cdc10 (Figure 1B,C,D). Since we measured integrated signal density, we considered that differences in cell size could indirectly alter signals between genotypes, but when we corrected for cell size the same changes were still apparent (Figure S1E and data not shown). Finally, Hsp104 BiFC signal was higher with Cdc10(D182N) (Figure 1C,D), recapitulating our published biochemical results (Johnson *et al*. 2015).

Encouragingly, many of the BiFC differences we saw linked directly to our previous functional studies. The prefoldin subunit Gim3 is required for efficient KADO of Cdc10(D182N), as Cdc10(D182N)-GFP localizes to septin rings in a subset of *gim3Δ* haploid cells co-expressing WT Cdc10 (Johnson *et al*. 2015). Ydj1 is also required for Cdc10(D182N) KADO and Ydj1 overexpression inhibits the proliferation of *cdc10(D182N)* cells (Johnson *et al*. 2015). Hsp104 is dispensable for septin KADO but Hsp104 overexpression inhibits the function of mutant septins subject to KADO (Johnson *et al*. 2015). We note that the decreases in Cdc10 interactions with Ydj1 and prefoldin subunits upon introduction of a mutation predicted to affect folding state are qualitatively similar to those observed with WT Cdc10 in cells cultured at 37°. These results are consistent with a model in which changes in the folding pathway of Cdc10(D182N) compared to WT Cdc10 correspond to changes in the identity and/or duration of the interactions made with cytosolic chaperones.

To ask if these observations with Cdc10 could be extended to another slow-folding mutant septin, we tested Cdc3(G365R), which we have previously shown is slower to incorporate into septin complexes even at a temperature permissive for function and in the absence of a WT allele of *CDC3* (Schaefer *et al*. 2016). We used a haploid strain carrying an untagged copy of the *cdc3(G365R)* allele at the *CDC3* locus and introduced a V_C_-tagged allele of *cdc3(G365R)* on a low-copy plasmid, or WT *CDC3* as a control. We then mated this strain with haploid strains carrying selected V_N_-tagged chaperones. This strategy allowed the tagged mutant septin to compete with the substoichiometric molecules of untagged WT Cdc3 and successfully localize to septin rings, as evidenced by bud neck fluorescence in cells co-expressing Hsp82-V_N_ (Figure 1E). That Cdc3(G365R)-V_C_ showed clear BiFC signal with Hsp82-V_N_ suggests that the decrease in Cdc10(D182N)–Hsp82 signal (Figure 1C,D) may be specific to that mutant septin. Examination of septin ring fluorescence in cells expressing V_N_-tagged Ydj1 revealed a decrease in BiFC signal for Cdc3(G365R)-V_C_ compared to Cdc3-V_C_ (Figure 1E), consistent with our results for Cdc10(D182N) and the decrease in Ydj1 interaction with WT Cdc3 observed at 37°. In summary, among the changes in Cdc10–chaperone interactions observed by BiFC upon introduction of a mutation predicted to slow Cdc10 folding are the two chaperones (Gim3 and Ydj1) that we have already shown to be functionally required for septin KADO, and the two chaperones (Hsp104 and Ydj1) for which we previously found genetic and biochemical evidence of increased physical associations with the mutant septin (Johnson *et al*. 2015). We conclude that BiFC is a useful tool to identify candidate chaperones whose expression can influence the function of misfolded mutant proteins.

### Co-expression of WT p53 fails to rescue the folding defects of the p53(V272M) mutant at high temperature

Having used mutant septins to establish BiFC as a useful tool, we wanted to identify differential chaperone interactions with mutant alleles of p53. However, our KADO model does not require chaperone interactions (Schaefer *et al*. 2016). Hence we first addressed other mechanisms that could prevent certain p53 mutants from acting dominantly to inhibit the function of the WT.

It was previously proposed (Dearth *et al*. 2007) that the reason some misfolded p53 mutants are recessive is because their proper folding is restored via interaction with properly-folded WT p53 molecules in the context of “mixed” hetero-tetramers. The V272M mutant adopts a non-native conformation in a temperature-sensitive manner (North *et al*. 2002; Blanden *et al*. 2020) and acts recessively when co-expressed with WT p53 in yeast cells harboring a *URA3* reporter gene containing p53 binding sites in the promoter region (Brachmann *et al*. 1996; Dearth *et al*. 2007) (see Figure 3). To ask if WT p53 is able to “chaperone” the proper folding of p53(V272M), we used a split-dihydrofolate reductase (split-DHFR) assay (Pittman *et al*. 2012). Here, p53 is fused between two fragments of DHFR and only folded p53 supports DHFR activity, manifested as proliferation by *dfr1Δ* yeast cells on medium lacking exogenous dTMP (Figure 2A). As shown in Figure 2B, *dfr1Δ* cells expressing a split-DHFR construct with p53(V272M) or p53(Y220C), another recessive and TS mutant (Dearth *et al*. 2007), barely proliferated at 37° on medium lacking dTMP. Equivalent constructs with WT or p53(R273H), a contact mutant that folds normally but cannot interact with DNA (Bullock *et al*. 1997), supported faster colony growth at all temperatures (Figure 2B). Co-expressing untagged WT p53 had no obvious effect when proliferation was assessed using colony growth on solid medium (Figure 2C). We also used a liquid culture growth assay and here observed a very slight effect of questionable significance (Figure 2D). Based on these data, we think folding rescue via interaction with WT p53 is unlikely to explain why the mutants do not act dominantly to interfere with WT function.

**Figure 2.**
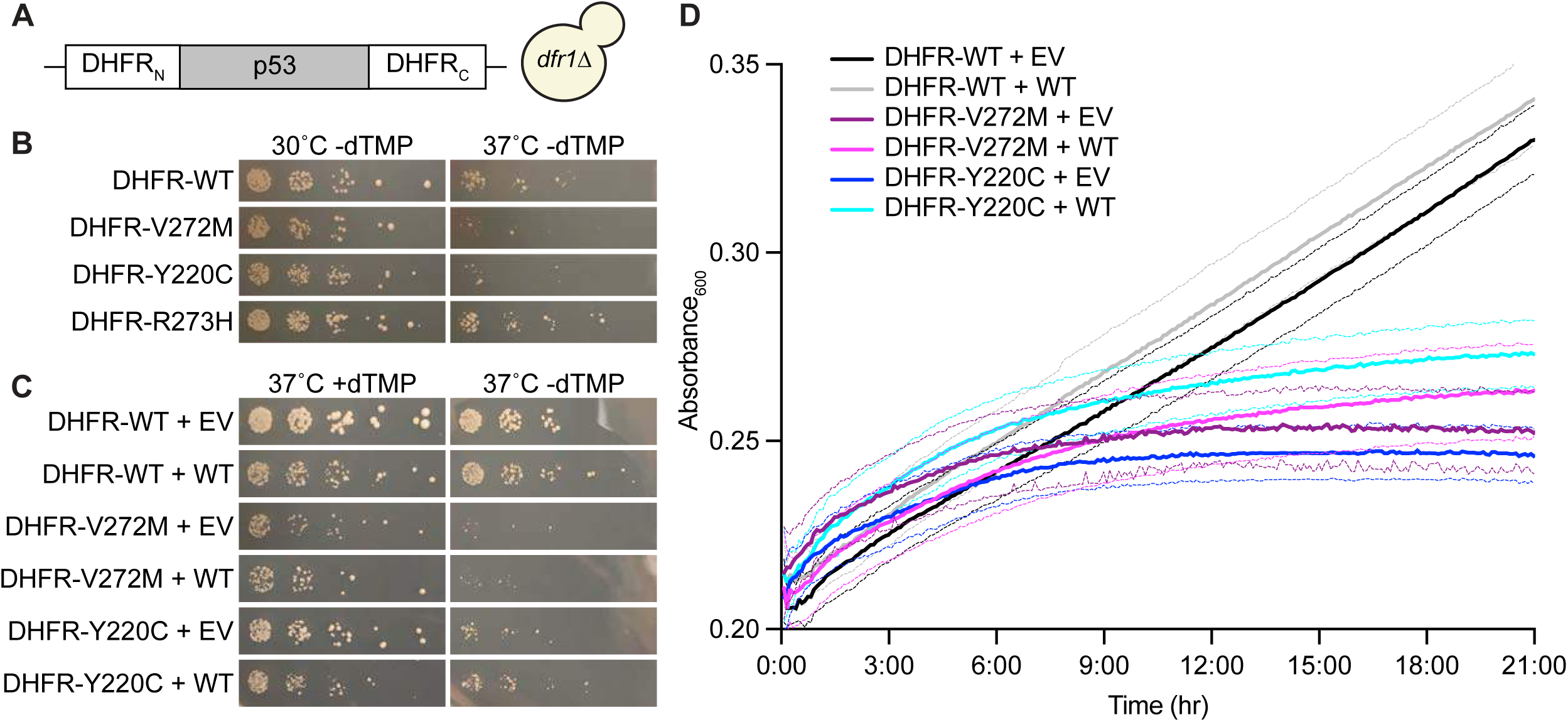
Co-expression of WT p53 fails to rescue the folding defects of the p53(V272M) mutant at high temperature. (A) Illustration of split dihydrofolate reductase (DHFR) p53 construct. TH5 cells are *Δdfr1* and proliferate only when exogenous dTMP is present or p53 folds well enough to reconstitute active DHFR. (B) Colony growth at 30° and 37° by fivefold serially diluted TH5 cells carrying plasmids encoding the indicated p53 split DHFR alleles spotted on solid synthetic medium selective for the plasmids and lacking dTMP. Plasmids were G00598, “DHFR-WT”; G00604, “DHFR-V272M”; G00599, “DHFR-Y220C”; G00605, “DHFR-R273H”. (C) As in (B), but with cells also carrying a plasmid encoding WT p53 (pLS76) or an empty vector (“EV”, pRS415) on solid medium selective for the plasmids and either containing “+” or lacking “-” dTMP at 37°. (D) As in (C) except cells were grown in liquid medium without dTMP at 37° in a flat-bottom 96- well plate. Optical density at 600 nm was recorded at 5-minute intervals over 21 hours for 6 replicate cultures per genotype and plotted as mean (thick line) and upper and lower bounds of the 95% confidence interval (thin lines).

**Figure 3.**
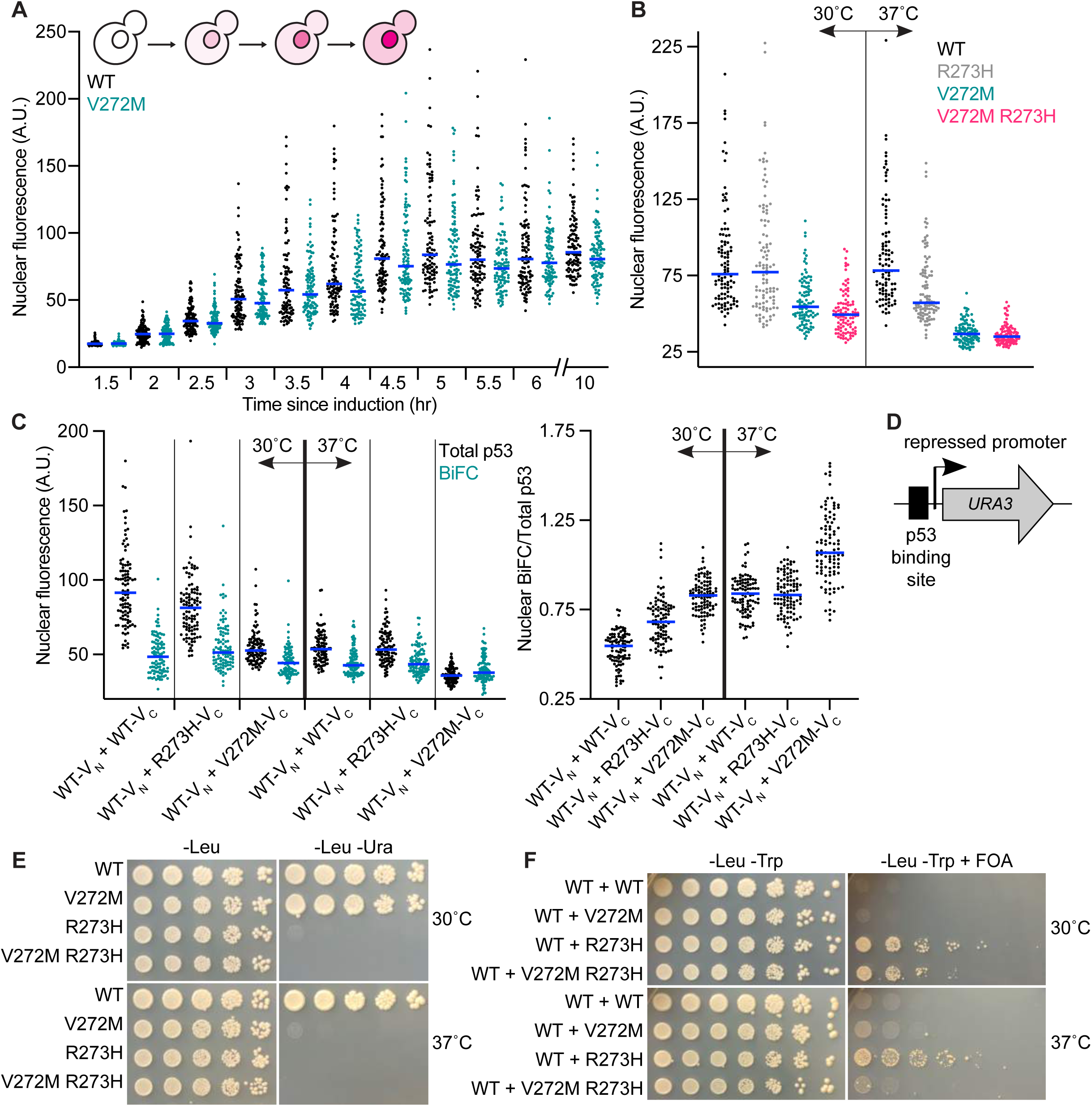
Effects of the V272M and R273H mutations on p53 nuclear localization, oligomerization, and function. (A) Nuclear mKate2 fluorescence in yJM3164 cells carrying a plasmid encoding *GAL1/10*-driven p53-mKate2-V_C_ (H3.29) or p53(V272M)- mKate2-V_C_ (pMAM67). Cells were imaged at the indicated time points following galactose induction at 24°. Blue lines denote median values. The schematic illustrates accumulation of nuclear and cytosolic p53 over time. (B) Steady-state nuclear mKate2 fluorescence of yJM3164 cells carrying plasmids encoding the indicated alleles of p53- mKate2-Venus and grown to mid-log phase at 30° or 37°. Blue lines denote median values. Vertical line separates 30° from 37°. Plasmids were pMAM85, “WT”; pMAM87, “R273H”; pMAM86, “V272M”; pMAM104, “V272M R273H”. (C) As in (B) but with cells of strain Y0134 co-expressing two p53 alleles fused to mKate2 and the indicated Venus fragments. Venus fluorescence here indicates physical interaction (BiFC). Left plot, total p53 (mKate2) and BiFC (Venus) nuclear fluorescence. Right plot, BiFC (Venus) nuclear signal normalized to total p53 (mKate2) nuclear signal. Blue lines denote median values. Vertical lines separate genotypes. Bold vertical lines separate 30° from 37°. Plasmids were pMAM98, “WT-V_N_”; pMAM82, “WT-V_C_”; pMAM84, “R273H-V_C_”; pMAM83, “V272M-V_C_”. (D) Schematic of p53 transcriptional reporter strains RBy33 and RBy159. Binding of properly folded p53 to a repressed promoter activates transcription of the *URA3* gene product, leading to colony growth on medium lacking uracil, or inhibition of growth on medium containing 5-fluoro-orotic acid (FOA). (E) Colony growth at 30° and 37° by RBy33 cells carrying plasmids encoding the indicated alleles of p53 on medium selective for the plasmids “-Leu” or selective for the plasmid and p53 reporter activity “-Leu -Ura”. Plasmids were pLS76, “WT”; pMAM69, “V272M”; pMAM79, “R273H”; pMAM77, “V272M R273H”. (F) Colony growth at 30° and 37° by diploid cells carrying plasmids encoding the indicated alleles of p53, together with WT p53 plasmid, on solid medium selective for the plasmids “-Leu -Trp” or selective for the plasmids and counter-selective for p53 reporter activity “-Leu -Trp + FOA”. Plasmids were pMAM78 and pLS76, “WT”; pMAM69, “V272M”; pMAM79, “R273H”; pMAM77, “V272M R273H”. Strains were diploids made by mating RBy33 cells carrying pLS76, pMAM69, pMAM79, or pMAM77 with RBy159 cells carrying pMAM78.

### High temperature inhibits p53(V272M) nuclear localization and co-assembly with WT p53

We next asked if kinetic delays during *de novo* synthesis and/or folding of p53(V272M) relative to WT p53 might prevent V272M mutant molecules from co-assembling with WT molecules into dysfunctional oligomers, akin to the KADO phenomenon we observed for septins. Oligomerization buries a nuclear export signal and ensures nuclear p53 localization (Stommel *et al*. 1999). Similar to how we monitored the kinetics of septin synthesis, folding and assembly by asking how long it takes a fluorescently-tagged septin to reach the septin ring at the bud neck (Schaefer *et al*. 2016), we monitored nuclear mKate2 signal after induction of mKate2-tagged p53 or p53(V272M). Since steady-state nuclear levels of WT p53 and p53(V272M) were equivalent at 24° (Figure 3A 10-hr timepoint and data not shown), we performed our experiments at this temperature. Our quantitative analysis demonstrated similar kinetics of nuclear accumulation for both WT and p53(V272M) (Figure 3A). Effect sizes with associated 95% confidence intervals for these and other quantifications of p53 fluorescence can be found in Figure S2. These results suggest that at low temperature mutant p53 is not significantly delayed relative to WT p53 in achieving the oligomerization-competent conformation required to reach the nucleus.

By contrast, nuclear localization of p53(V272M)-mKate2 was reduced at 30° and greatly reduced at 37° (Figure 3B). Temperature had a much smaller effect, if any, on nuclear levels of WT p53 and the “contact” mutant R273H (Figure 3B). These data point to a failure in nuclear localization as a major cause of p53(V272M) dysfunction at high temperature. The inability of V272M to dominantly interfere with WT p53 when the two alleles are co-expressed may reflect a reduced ability of V272M to co-assemble with WT molecules into “mixed” oligomers wherein the mutant perturbs WT function. If WT p53 interacts poorly with p53(V272M) at 37° that would additionally explain the results of our split-DHFR experiments.

We next used BiFC to test for p53–p53 interactions in living cells, using p53-mKate2-V_C_ and p53-mKate2-V_N_ constructs previously validated in mammalian cells (Gaglia and Lahav 2014). We quantified nuclear signal for total p53 (mKate2 fluorescence) and for p53–p53 BiFC (Venus fluorescence) at 30° and 37°. Whereas total nuclear p53 and BiFC signals indicative of p53–p53 interactions were indistinguishable for WT–WT and WT–R273H at 37°, for WT–V272M both total nuclear p53 and BiFC signals were reduced (Figure 3C). Since WT p53 also contributed to nuclear signal, this means that at 37° co-expression of the V272M mutant partially inhibited the nuclear localization of WT p53. By normalizing nuclear BiFC signal to total nuclear p53 signal, we found that the relative proportion of nuclear p53 present as “mixed” oligomers was much higher for WT–V272M than WT–WT or WT–R273H (Figure 3C). These data indicate that at 37° only a small number of V272M molecules are able to interact with WT p53 and remain the nucleus, supporting a model in which at high temperature V272M mostly fails to fold into an oligomerization-competent conformation and thus cannot inhibit WT p53 function.

### The V272M mutation converts the dominant R273H mutant to a recessive form

If p53(V272M) is recessive because it misfolds and fails to interact with WT p53, and p53(R273H) is dominant because it interacts with WT p53 to form dysfunctional tetramers, then introducing the V272M mutation should sequester the R273H mutant in the cytosol and render it recessive. Indeed, quantification of nuclear fluorescence revealed that nuclear localization of p53(V272M R273H) was diminished at 30° and greatly diminished at 37°, patterns indistinguishable from that of p53(V272M) (Figure 3B). Like p53(R273H), when expressed alone p53(V272M R273H) was transcriptionally inactive at all temperatures tested (Figure 3D,E). At 30°, both p53(R273H) and p53(V272M R273H) dominantly inhibited WT p53 function (Figure 3F). However, at 37° p53(V272M R273H) was unable to inhibit WT function (Figure 3F), consistent with our prediction. These data point to temperature-induced misfolding and cytosolic sequestration as the major mechanisms by which the V272M mutation perturbs oligomerization and compromises p53 function.

### Mutations to Leu344 do not specifically perturb p53 oligomerization in the context of expression in yeast

Our findings demonstrating the effects of V272M on p53 oligomerization led us to ask when during p53 oligomerization the defects occurred. In previous studies, including during cell-free translation *in vitro* (Nicholls *et al*. 2002), p53 dimerizes co-translationally (Gaglia and Lahav 2014) such that tetramers assemble from dimers composed of two monomers encoded by the same mRNA. Thus distinct alleles do not “mix” in the dimer form but rather in the tetramer form only. Mutants that block oligomerization before the dimerization (L344P) or tetramerization (L344A) step are useful tools for examining co-translational dimerization, particularly when combined with BiFC: the L344P and L344A mutations eliminate the majority of p53 BiFC signal in mammalian cells co-expressing WT p53-V_C_ and p53-V_N_ (Gaglia and Lahav 2014) as illustrated in Figure 4A.

**Figure 4.**
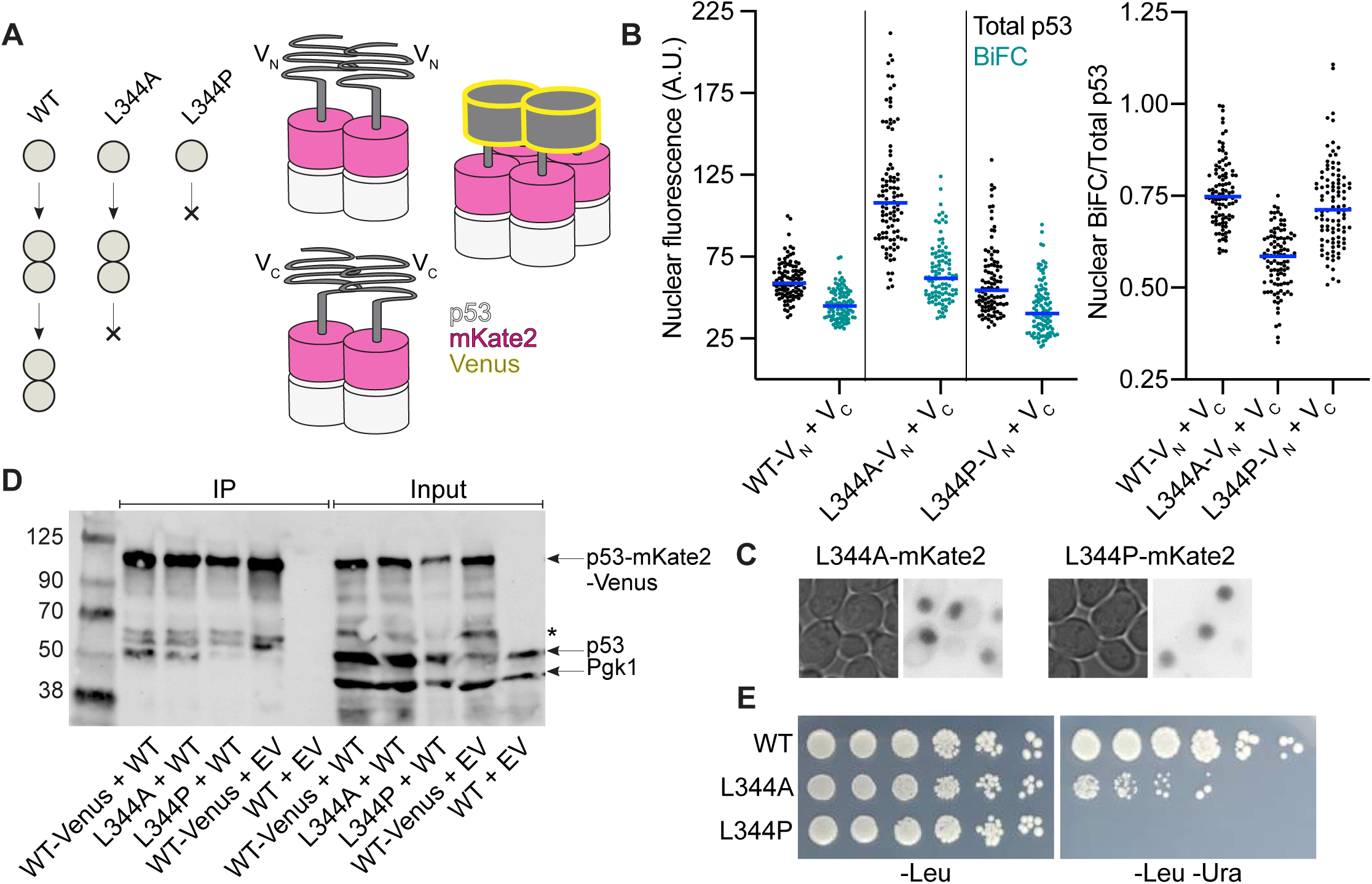
Mutations to Leu344 do not specifically perturb p53 oligomerization in the context of expression in yeast. (A) Schematic of oligomerization-disrupting mutants and expected p53 BiFC results. Left, L344 mutations destabilize higher-order p53 oligomerization. Alanine substitution yields dimers but not tetramers and proline substitution yields monomers only. Right, co-expression with co-translational dimerization of p53(L344A)-mKate2-V_N_ and p53(L344A)-mKate2-V_C_ results in V_N_ + V_N_ and V_C_ + V_C_ homodimers, neither of which reconstitute Venus. Venus fluorescence is only observed upon tetramer formation. Adapted from (Gaglia and Lahav 2014). (B) Total p53 (mKate2) and BiFC (Venus) nuclear fluorescence (left plot) as well as BiFC (Venus) nuclear signal normalized to total p53 (mKate2) nuclear signal (right plot) of Y0134 cells carrying plasmids encoding the indicated p53 BiFC alleles. Cells were grown to mid-log phase at 37° prior to imaging. Blue lines denote median values. Vertical lines separate genotypes. Plasmids were pMAM98, “WT-VN”; pMAM82, “WT- V_C_”; pMAM99, “L344A-V_N_”; pMAM91, “L344A-V_C_”; pMAM100, “L344P-V_N_”; pMAM93, “L344P-V_C_”. (C) Representative micrographs showing nuclear localization of p53(L344A) and p53(L344P) (mKate2 fluorescence, inverted to facilitate viewing) in BiFC experiments. Cells were grown to mid-log phase at 37° prior to imaging. Plasmids were pMAM99, “L344A-mKate2-V_N_”; pMAM91, “L344A-mKate2-V_C_”; pMAM100, “L344P- mKate2-V_N_”; pMAM93, “L344P-mKate2-V_C_”. (D) JTY4202 cells carrying plasmids encoding the indicated alleles of p53 were grown to mid-log phase at 30° and lysed. Venus-tagged proteins were immunoprecipitated with Venus-binding nanobodies and separated by SDS–PAGE. After electrophoretic transfer, the membrane was probed for p53 and the cytosolic protein Pgk1. Asterisk, proteolytic degradation products derived from p53-mKate2-Venus. pRS415 and pRS314, “EV”; pMAM78, “WT”; pMAM85, “WT-Venus”; pMAM105, “L344A-Venus”; pMAM106, “L344P-Venus”. EV, empty vector. (E) Colony growth at 30° by RBy33 cells carrying plasmids encoding the indicated alleles of p53 on medium selective for the plasmids (“-Leu”) or selective for the plasmid and p53 reporter activity (“-Leu -Ura”). Plasmids were pLS76, “WT”; pMAM114, “L344A”; pMAM115, “L344P”.

To our surprise, in yeast we observed robust BiFC signal between p53(L344A)- mKate2-V_N_ and p53(L344A)-mKate2-V_C_ and, most unexpectedly, between p53(L344P)- mKate2-V_N_ and p53(L344P)-mKate2-V_C_ (Figure 4B). Leu 344 lies within a nuclear export signal that is normally buried in the context of a tetramer (Stommel *et al*. 1999). Presumably because the export signal they expose is non-functional, p53(L344P) and p53(L344A) both localize to the nucleus in mammalian cells, albeit in monomeric and dimeric form, respectively (Gaglia *et al*. 2013). L344A and L244P also localized primarily to the nucleus in our yeast cells (Figure 4C). In fact, when we quantified nuclear mKate2 fluorescence, we found that the L344A mutant showed even higher total nuclear signal than WT (Figure 4B). Accordingly, normalizing the BiFC signal to total nuclear p53 signal for each cell resulted in BiFC/total p53 ratios for L344A that were somewhat less than WT (Figure 4B). However, the “monomeric” L344P mutant ratios and the raw BiFC signals for both mutants were equivalent to WT (Figure 4B). We interpret these data as evidence that in yeast cells p53 oligomerization is less sensitive to mutations known to disrupt oligomerization in mammalian cells.

To independently test for co-translational dimerization of WT p53 expressed in yeast cells, we immunoprecipitated p53-mKate2-Venus from lysates of cells co-expressing untagged p53. Whereas in mammalian cells two differentially tagged forms co-precipitate only when residue 344 is Leu (Gaglia and Lahav 2014), we saw evidence of “mixed” complexes (i.e., co-precipitation of untagged WT p53) even in the presence of the L344A mutation (Figure 4D). The L344P mutation for the most part blocked co-IP (Figure 4D), and in the *URA3* reporter strain p53(L344P) was functionally dead (Figure 4E). We thus interpret our BiFC data as evidence that the complementation event itself stabilizes otherwise weak interactions between p53(L344P) molecules. In the same assay, p53(L344A) displayed moderate transcriptional activation (Figure 4E). This result is consistent with published yeast reporter experiments (Jordan *et al*. 2008) and the persistent (albeit reduced) ability of this mutant to bind DNA *in vitro* (McLure and Lee 1998). Because in our yeast cells the L344A and L344P mutations were not reliable disruptors of dimerization or tetramerization, our experiments were unable to distinguish between co-and post-translational mechanisms of dimerization or determine the point at which oligomerization fails in the V272M mutant.

### Cytosolic chaperone-p53 interactions in living cells

Given the evidence that misfolding of p53 mutants is responsible for dysfunction at high temperatures (reviewed in (Bullock and Fersht 2001)) and drives prolonged associations with at least three kinds of cytosolic chaperones (Sepehrnia *et al*. 1996; Blagosklonny *et al*. 1996; Singh *et al*. 2010; Wiech *et al*. 2012), we used our BiFC approach to systematically identify p53–chaperone interactions. We predicted that the conformationally unstable p53(V272M) would have altered chaperone interactions relative to WT p53 or the “contact” mutant p53(R273H). We used most of the same V_N_-tagged chaperones as with septins, paired with the p53-mKate2-V_C_ fusion from the p53–p53 BiFC experiments described above. (Note that the V_C_ tag here is slightly larger than the V_C_ tag used with septins.)

We identified interactions between p53 and numerous cytosolic chaperones (Figure 5A, Figure S3A, and summarized in Table 5). To account for differential proteasomal degradation of p53 mutants (Singh *et al*. 2010), we used the mKate2 tag to normalize the p53-chaperone BiFC signal (Venus YFP) relative to total p53 signal (mKate2), which revealed increased interactions between p53(V272M) and many chaperones relative to WT p53 and p53(R273H) (Figure 5B). Several chaperones – Hsp82, Ssb1, Sis1, and Ydj1 – demonstrated increases in p53(V272M) interaction that were obvious even before normalization (raw values in Figure S3B).

**Figure 5.**
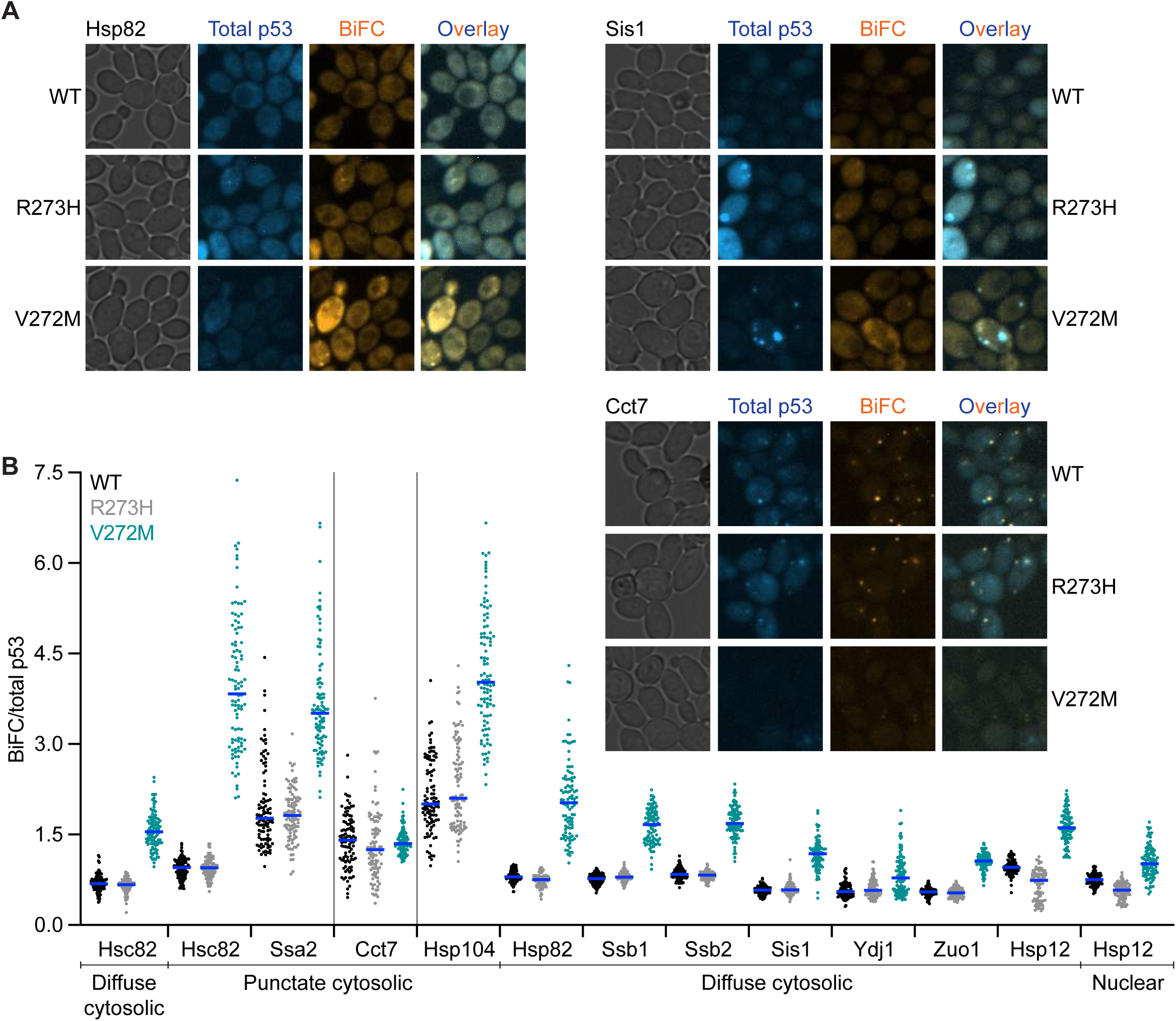
Cytosolic chaperone-p53 interactions in living cells. (A) Representative micrographs of p53–chaperone BiFC interactions in cells cultured at 30° and expressing Hsp82-V_N_, Sis1-V_N_, or Cct7-V_N_ and carrying plasmids encoding the indicated p53- mKate2-V_C_ alleles. Total p53 (mKate2) fluorescence images were false-colored blue and BiFC (Venus) fluorescence images were false-colored orange. Prior to overlay, brightness and contrast were adjusted equivalently for each set of images. Plasmids were pMAM110, “WT”, pMAM112, “R273H”; pMAM111, “V272M”. Strains were diploids made by mating BY4742 cells carrying the plasmids with H00398, H06680, or H00383. (B) BiFC (Venus) signal normalized to total p53 (mKate2) signal in cells cultured at 30° carrying plasmids encoding the indicated p53-mKate2-V_C_ alleles together with the indicated V_N_-tagged chaperone. Blue lines denote median values. Vertical lines group the samples imaged with the same settings (LED intensity and exposure time). The subcellular location where the BiFC interaction was quantified is indicated below. Plasmids were pMAM110, “WT”, pMAM112, “R273H”; pMAM111, “V272M”. Strains were diploids made by mating BY4742 cells carrying the plasmids with H00399, “Hsc82”; H00388, “Ssa2”; H00383, “Cct7”; H00384, “Hsp104”; H00398, “Hsp82”; H00395, “Ssb1”; H00394, “Ssb2”; H06680, “Sis1”; H00385, “Ydj1”; H00396, “Zuo1”; or H06684, “Hsp12”.

**Table 5.**
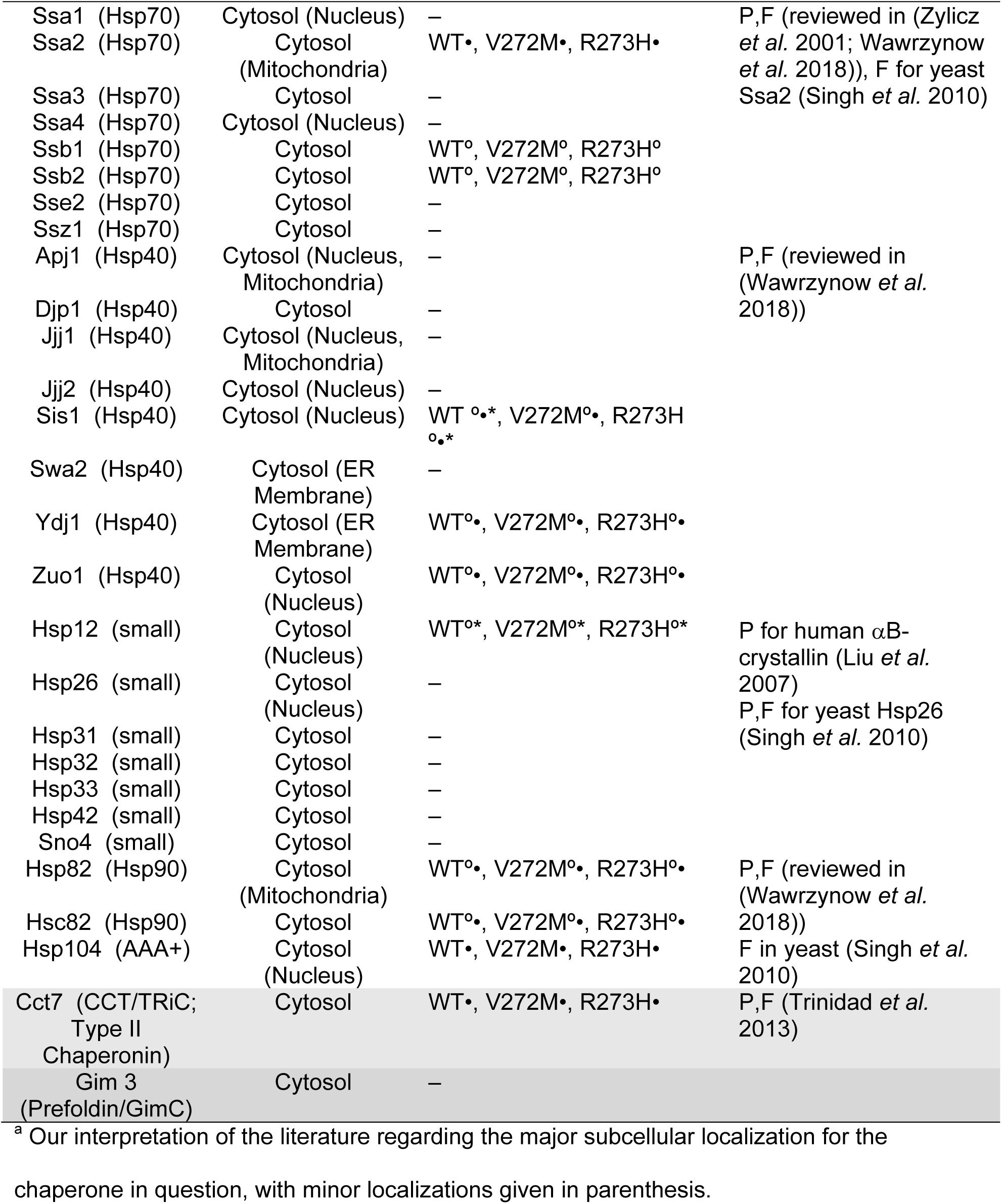
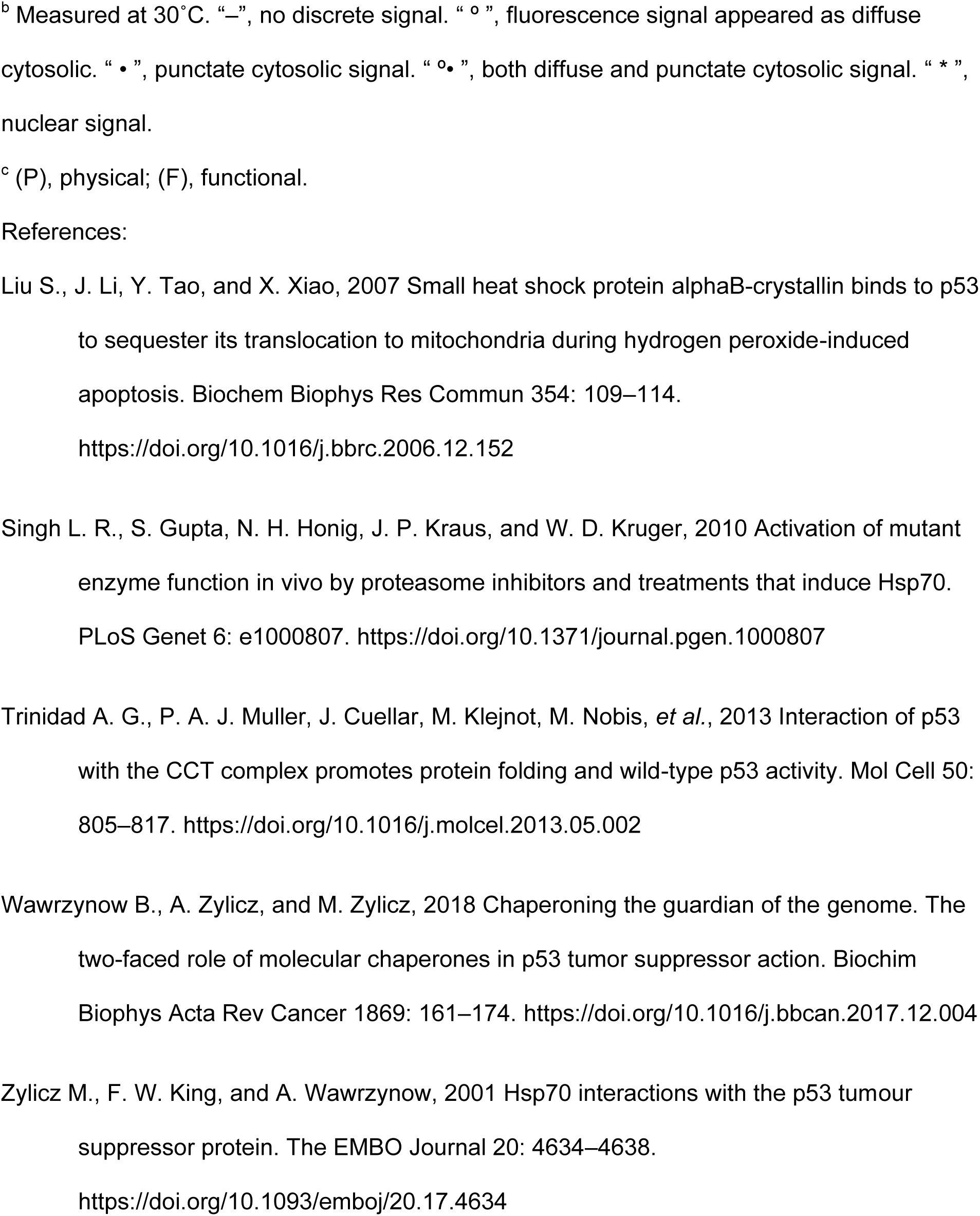
p53–chaperone interactions assayed by BiFC

With the exception of Cct7 and Hsp12, and Hsp104, which generated decreased and increased BiFC signal with p53(R273H) compared to WT, respectively, most chaperones interacted in similar ways with p53(R273H) and WT p53, whereas p53(V272M) displayed multiple distinct interactions (Figure 5B and Figure S3B). p53(R273H) specifically abolishes contact with DNA without global conformational consequences (Ang *et al*. 2006). On the other hand, p53(V272M) causes misfolding and conformational instability (North *et al*. 2002; Blanden *et al*. 2020). These observations are consistent with a model in which chaperones recognize specific, exposed sequences in p53 that are largely buried in the native fold. The V272M mutation results in prolonged exposure of these sequences and thus more opportunities for cytosolic chaperone engagement.

In some cases, we also saw differences in BiFC signal localization. At 30°, the temperature at which we cultured cells prior to microscopic examination, the mKate2- V_C_-tagged p53 alleles differed in their localization even when no V_N_-tagged chaperone was expressed. WT p53 and the R273H mutant were seen in both the cytosol and the nucleus, whereas the V272M mutant was found almost exclusively in the cytosol (Figure S3A, “No BiFC Control”). These observations agree with prior cell fractionation experiments in mammalian cells, in which WT p53 and p53(R273H) were found almost exclusively in the nuclear fraction, regardless of temperature (Zhang *et al*. 1994) whereas p53(V272M) showed a distinct shift toward the cytosolic fraction even at low temperatures (Dearth *et al*. 2007). Most p53–chaperone BiFC signals were in the cytosol, either diffusely, as puncta, or a mixture of both signal types (Table 5). Hsp70 chaperones Ssb1 and Ssb2 generated diffuse cytosolic signal whereas Ssa2, Cct7, and Hsp104 generated exclusively punctate signals (Table 5, Figure 5A, Figure S3A). Multiple p53–Cct7 puncta were seen in some cells (Figure 5A), clearly distinguishing them from septin–Cct7 puncta. Chaperones producing a mixture of both diffuse and punctate cytosolic signal include the Hsp90 isoforms Hsp82 and Hsc82 (with brighter and more numerous puncta for Hsc82) and the Hsp40 family members Sis1, Ydj1, and Zuo1 (Table 5, Figure 5A, Figure S3A). Sis1 and Hsp12 were unique in producing nuclear signal in addition to cytosolic fluorescence (Figure 5A, Figure S3A). Sis1 generated nuclear signal for WT and p53(R273) but essentially none for p53(V272M) (Figure 5A).

By monitoring the mKate2 signal, we found several cases in which co-expression of a V_N_-tagged chaperone altered the localization of mKate2-V_C_-tagged p53 (Figure 5A and Figure S3A). These observations are consistent with a model in which Venus reconstitution trapped the p53 molecules in the location where the chaperone is normally found. Nuclear mKate2 signal for p53(V272M) in cells co-expressing Hsp12-V_N_ is a particularly striking example (Figure S3A). In all cases, the Venus signal co-localized with the p53 mKate2 signal in the “new” location, further arguing that BiFC events were responsible (Figure 5A and Figure S3A). We interpret these data as evidence that, just as in septin–chaperone BiFC, making permanent an otherwise transient interaction incites a localization “tug-of-war”. For most septin–chaperone interactions, the septin wins; for all p53–chaperone interactions, the chaperone wins. The results described above point to a number of chaperones as compelling candidates to influence the function of mutant alleles of p53.

### Overexpression of specific chaperones inhibits p53(V272M) function

Differences in chaperone engagement might explain functional differences between p53(V272M) and WT p53 if a chaperone sequesters the mutant protein in a non-functional state. We asked if the function of p53 could be aggravated by chaperone overexpression at a permissive temperature, focusing on the chaperones identified by our BiFC experiments. Plasmids encoding chaperones under control of a galactose-inducible promoter were introduced into a diploid *URA3* reporter strain also carrying a plasmid encoding untagged WT p53 or p53(V272M) under control of a constitutive promoter. Various cytosolic chaperones impeded the transcriptional function of p53(V272M) and, to a generally lesser degree, WT p53 (Figure 6A). Exceptions included Hsf1ΔN, a constitutively active version of the major heat shock transcriptional regulator, where WT p53 and p53(V272M) behaved similarly (Park *et al*. 2006). By contrast, Hsf1ΔN, like expression of specific individual chaperones, inhibits the proliferation of cells with the *cdc10(D182N)* septin mutation (Johnson *et al*. 2015). Hsp90 isoforms Hsp82 and Hsc82 slightly but equivalently impeded mutant p53(V272M) relative to WT p53 although the intensity of signal and number of puncta were increased for endogenous levels of Hsc82 in BiFC experiments. Both Hsp90 isoforms were expressed at similar levels, as assessed by immunoblotting (Figure 6B).

**Figure 6.**
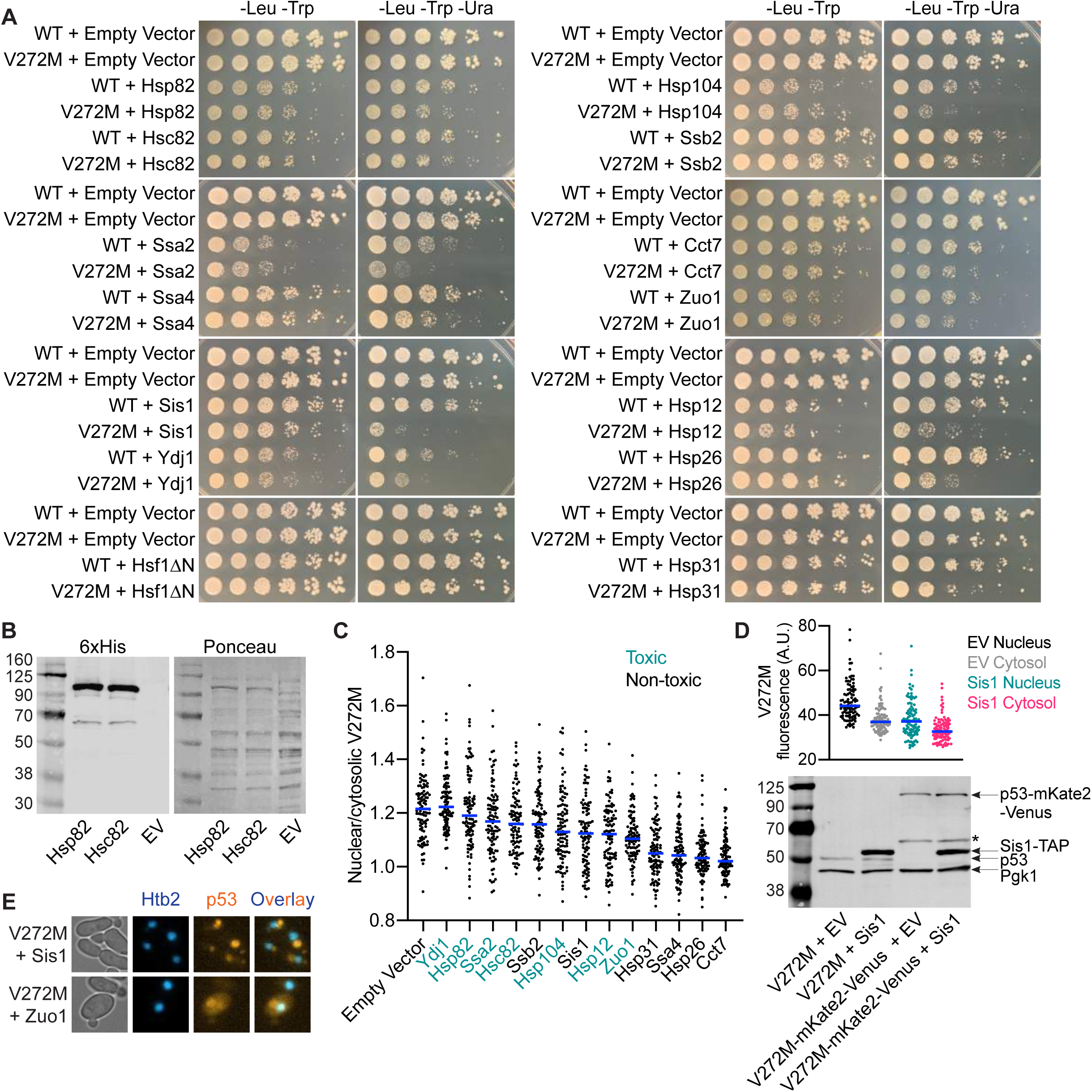
Overexpression of specific chaperones inhibits p53(V272M) function. (A) Colony growth at 30° of fivefold serially diluted diploid cells carrying plasmids encoding the indicated alleles of p53, together with the indicated *GAL1/10*-driven chaperone plasmid or an empty vector, on solid galactose medium selective for the plasmids “-Leu -Trp” or selective for the plasmids and p53 reporter activity “-Leu -Trp -Ura”. Plasmids were pLS76, “WT”; pMAM69, “V272M”; pRS424 and pRS314, “Empty Vector”; G00630, “Hsp82”; G00799, “Hsc82”; G00633, “Ssa2”; G00789, “Ssa4”; G00632, “Sis1”; G00353, “Ydj1”; G00061, “Hsf1ΔN”; G00631, “Hsp104”; G00634, “Ssb2”; G00620, “Cct7”; G00635, “Zuo1”; G00627, “Hsp12”; G00628, “Hsp26”; G00629, “Hsp31”. Strains were diploids made by mating RBy33 cells carrying pLS76 or pMAM69 with RBy159 cells carrying the *GAL1/10*-driven chaperone plasmid or empty vector. (B) SDS-PAGE and immunoblot analysis of total protein extracts from diploid cells carrying a plasmid encoding p53(V272M) (pMAM69) together with a *GAL1/10*-driven Hsp82 plasmid (G00630), a *GAL1/10*-driven Hsc82 plasmid (G00799), or an empty vector (pRS424, “EV”) grown to mid-log phase at 30° in medium with galactose to induce overexpression. The membrane was first probed for hexahistidine-tagged Hsp82 and Hsc82 and subsequently stained with Ponceau. (C) Ratio of nuclear to cytosolic mKate2 signal in diploid cells carrying a plasmid encoding p53(V272M)-mKate2-Venus together with the indicated *GAL1/10*-driven chaperone plasmid or an empty vector. Cells were grown to mid-log phase at 30° with galactose to overexpress chaperones prior to imaging. “Toxic” denotes chaperones whose overexpression renders p53 inhibitory to colony growth independent of p53 function. Blue lines denote median values. Plasmids were pMAM86, “p53(V272M)”; pRS424, “Empty Vector”; G00353, “Ydj1”; G00630, “Hsp82”; G00633, “Ssa2”; G00799, “Hsc82”; G00634, “Ssb2”; G00631, “Hsp104”; G00632, “Sis1”; G00627, “Hsp12”; G00635, “Zuo1”; G00629, “Hsp31”; G00789, “Ssa4”; G00628, “Hsp26”; G00620, “Cct7”. Strains were diploids made by mating yJM3164 cells carrying pMAM86 with yJM1838 cells carrying the indicated *GAL1/10*-driven chaperone plasmid or empty vector. (D) Top, nuclear and cytosolic mKate2 fluorescence in diploid cells carrying a plasmid encoding p53(V272M)-mKate2-Venus (pMAM86) and an empty vector (“EV”, pRS424) or a plasmid encoding tandem affinity purification-(TAP-)tagged Sis1 (G00632). Cells were grown to mid-log phase at 30° with galactose to overexpress Sis1 prior to imaging. A single outlier (Y=115) was omitted from the EV Nucleus dataset to facilitate viewing on the Y-axis. Blue lines denote median values. Strains were diploids made by mating yJM3164 cells carrying pMAM86 to yJM1838 cells carrying G00632 or pRS424. Bottom, as in (B) but with diploid cells carrying plasmids encoding the indicated alleles of p53 together with the Sis1-TAP plasmid or an empty vector, as in (A) or the plot above, and probed for p53, the cytosolic protein Pgk1, and hexahistidine-tagged Sis1-TAP. Asterisk, proteolytic degradation products derived from p53-mKate2- Venus. (E) Representative micrographs of vacuolar mKate2 signal in diploid cells expressing p53(V272M)-mKate2-Venus (pMAM86) and *GAL1/10*-driven Sis1 (G00632) or *GAL1/10*-driven Zuo1 (G00616), prepared as in (C). Nucleus (Htb2-mNeon) fluorescence images were false-colored blue and p53 (mKate2) fluorescence images were false-colored orange. Prior to overlay, brightness and contrast were adjusted equivalently for each set of images. Strain was a diploid made by mating yJM3164 cells carrying pMAM86 to yJM1838 cells carrying G00632 or G00616.

Since the transcriptional activation function we monitor here requires nuclear p53, one way chaperone engagement might inhibit p53(V272M) function is via cytosolic sequestration. We quantified nuclear versus cytosolic p53(V272M)-mKate2-Venus signal in micrographs and found that in most cases chaperone overexpression decreased the nuclear/cytosolic ratio of p53(V272M), with the exception of Ydj1 where the ratio was preserved (Figure 6C). Notably, in this imaging-based approach, cytosolic p53(V272M) levels sometimes also decreased along with nuclear levels relative to empty vector (Figure S4A,B), as illustrated for Sis1 (Figure 6D). However, when we tested Sis1 overexpression by immunoblotting, we saw no clear difference in total p53(V272M) levels relative to the empty vector control (Figure 6D). Our imaging-based quantification may underestimate total cytosolic levels, since we excluded vacuolar p53 signal seen in some chaperone-overexpressing cells (e.g., Sis1 and Zuo1, Figure 6E). We interpret these data as evidence that overexpression of specific chaperones sequesters p53(V272M) outside the nucleus and impedes transcriptional activation.

Overexpression of some chaperones inhibited colony growth in general, even when we did not select for p53 function, and this effect was independent of the p53 allele expressed (Figure 6A). Overexpression of these same chaperones in cells not co-expressing p53 has no such effect (Sopko *et al*. 2006; Johnson *et al*. 2015). High-level p53 expression is toxic to yeast and is thought to involve p53 binding to endogenous yeast genes and thereby perturbing essential gene expression (Inga and Resnick 2001; Leão *et al*. 2013; Park *et al*. 2021). We noticed that the chaperones with the strongest effect on non-selective growth (i.e., those that induced the most p53 “toxicity”) had the most p53(V272M) in the nucleus (Figure 6A,C and Figure S4A,B). Conversely, the chaperones that specifically inhibited p53(V272M) transcriptional activation function and most strongly decreased the levels of p53(V272M) in the nucleus (Hsp31, Ssa4, Hsp26, and Cct7) had little to no effect on non-selective growth (Figure 6A,C and Figure S4A,B). While the details of the underlying mechanism are beyond the scope of this work, these results support the idea that p53 toxicity in yeast requires nuclear localization. However, our findings further suggest that canonical transcriptional activation function is dispensable for toxicity, and that whatever function is responsible can be modulated by chaperone expression.

### Cytosolic compartmentalization of p53 modulates recessive behavior

If association with a specific chaperone mediates cytosolic sequestration, then inhibiting the ability of that chaperone to bind p53 should allow more p53 to reach the nucleus. Indeed, when we inhibited Hsp90 function using radicicol (Sharma and Masison 2011), nuclear levels and nuclear/cytosolic ratios increased for WT p53 and all variants (Figure S4C,D and Figure 7A). As assessed by immunoblotting, total p53 levels were not detectably changed between DMSO- and radicicol-treated cells (Figure 7B). We also observed vacuolar p53 signal in cells treated with radicicol but not with DMSO (Figure 7C), reminiscent of chaperone-overexpressing cells, and this vacuolar signal was not included in our quantification of cytosolic p53 (Figure S4C,D). These findings are consistent with other studies with mammalian cells in which pharmacological Hsp90 inhibition increased nuclear localization of a different p53 mutant (Whitesell *et al*. 1998). Notably, in that study Hsp90 inhibition did not improve the function of the mutant p53 (Whitesell *et al*. 1998). We reasoned that the liberation of non-functional mutant p53 from chaperone sequestration might increase the ability of the V272M mutant to interact with WT p53 and dominantly interfere with p53 function. Indeed, activation of the *URA3* reporter gene by WT p53 was reduced by the co-expression of p53(V272M) only in the presence of radicicol, as evidenced by a very slight increase in colony growth on FOA-containing medium (Figure 7D). Interestingly, radicicol also increased FOA growth in the absence of any co-expressed p53 mutant (Figure 7D). Considering that even WT p53 is conformationally labile and often samples non-functional conformations (Hainaut *et al*. 1995; Cañadillas *et al*. 2006), this result might suggests that Hsp90 inhibition allows molecules of WT p53 in non-functional conformations to translocate to the nucleus, where they cannot activate transcription but may dominantly interfere with the function of any properly-folded molecules.

**Figure 7.**
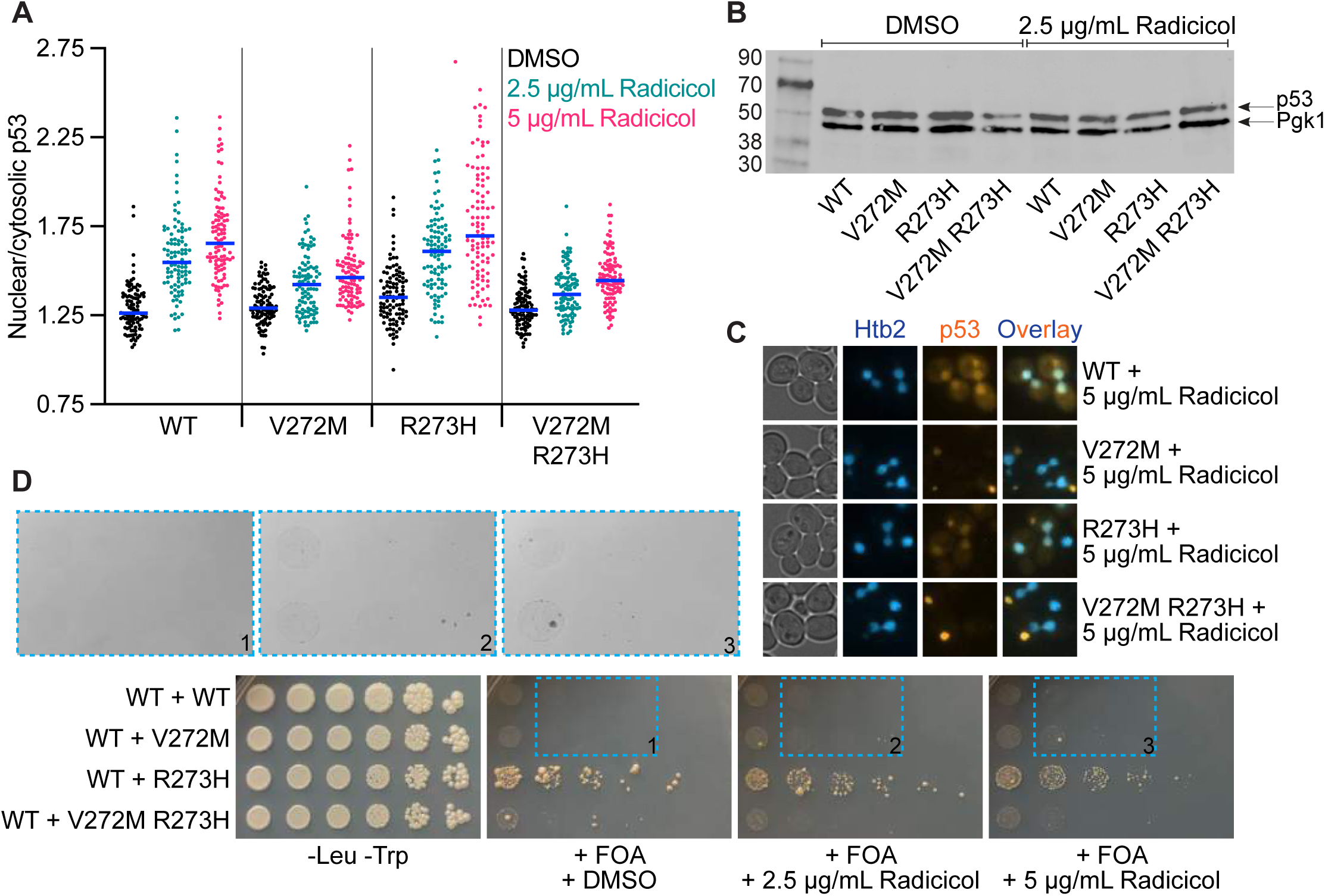
Cytosolic compartmentalization of p53 modulates recessive behavior. (A) Ratio of nuclear to cytosolic mKate2 signal in yJM3164 cells carrying plasmids encoding the indicated alleles of p53-mKate2-Venus. Cells were grown to mid-log phase at 37° with 2.5 µg/mL radicicol or equivalent volume of DMSO prior to imaging. Blue lines denote median values. Vertical lines separate genotypes. Plasmids were pMAM85, “WT”; pMAM86, “V272M”; pMAM87, “R273H”; pMAM104, “V272M R273H”. (B) RBy33 cells carrying plasmids encoding the indicated alleles of p53 were grown to mid-log phase at 30° and lysed. Proteins were separated by SDS–PAGE. After electrophoretic transfer, the membrane was probed for p53 and the cytosolic protein Pgk1. Plasmids were pLS76, “WT”; pMAM69, “V272M”; pMAM79, “R273H”; pMAM77, “V272M R273H”. (C) Representative micrographs of vacuolar mKate2 signal in yJM3164 cells carrying plasmids encoding the indicated alleles of p53-mKate2-Venus and treated with 5 µg/mL radicicol, prepared as in (A). Nucleus (Htb2-mNeon) fluorescence images were false-colored blue and p53 (mKate2) fluorescence images were false-colored orange. Prior to overlay, brightness and contrast were adjusted equivalently for each set of images. Plasmids were pMAM85, “WT”; pMAM86, “V272M”; pMAM87, “R273H”; pMAM104, “V272M R273H”. (D) Colony growth at 37° of diploid cells carrying plasmids encoding the indicated alleles of p53, together with WT p53 plasmid, on solid medium selective for the plasmids “-Leu -Trp” or selective for the plasmids and counter-selective for p53 reporter activity “-Leu -Trp + FOA” and containing the indicated concentration of radicicol or equivalent volume of DMSO. Magnified images were inverted and adjusted for brightness and contrast in an equivalent manner. Plasmids were pLS76, “WT”; pMAM69, “V272M”; pMAM79, “R273H”; pMAM77, “V272M R273H”. Strains were diploids made by mating RBy33 cells carrying pLS76, pMAM69, pMAM79, or pMAM77 with RBy159 cells carrying pMAM78.

## DISCUSSION

Chaperone binding to a misfolded protein is widely assumed to either promote proper folding or protect cells from adverse consequences of irreversibly misfolded proteins with no chance of ever reaching a functional state. Accordingly, chaperone upregulation or overexpression should rescue mutant protein function. However, there are some clear examples in which chaperones “inappropriately” sequester otherwise functional proteins, such as the ΔF508 mutant allele of CFTR, the most common mutation causing human cystic fibrosis (Amaral 2006; Balch *et al*. 2011; Lopes-Pacheco *et al*. 2015). Conversely, some mutant proteins can fold well enough to hetero-oligomerize with WT partner proteins but carry substitutions that “poison” the function of the hetero-oligomer. Here, chaperone sequestration can render recessive an otherwise dominant-negative mutant. We aimed to developed tools for identifying chaperones that differentially engage misfolding-prone mutant proteins, and use them to test if chaperone association affects the function and dominant-negative effect of a misfolding-prone mutant of p53.

Our BiFC approach identified differential chaperone interactions with mutant septins that matched results from our prior functional studies (Johnson *et al*. 2015). Other results provide further validation of BiFC as a useful tool. Only two of 11 intended negative controls, Sec63 and Pam18, generated detectable septin BiFC signal (Table 4). The Hsp40 family member Sec63 is normally part of a membrane-associated complex that mediates both co- and post-translational translocation of polypeptides across the ER membrane (Ng and Walter 1994). Sec63 co-purifies with dozens of proteins, many of which are cytosolic (Krogan *et al*. 2006). We suspect that in its post-translational role, Sec63 could transiently associate with many cytosolic proteins (including septins), only some of which are ultimately threaded into the ER lumen. Consistent with this interpretation, Sec63-V_C_ interacts with “free” V_N_ (Kim *et al*. 2019). Hence Sec63 is probably not a good negative control. The mitochondrially-localized chaperone Pam18 interacted only with Cdc3 and only at 37° (Table 4). While interactions between Cdc12 and Pam18 and between Cdc3 and Hsp78, another mitochondrial chaperone, were reported in high-throughput genetic (Davierwala *et al*. 2005) and two-hybrid (Wang *et al*. 2012) screens, we doubt that these interactions are biologically meaningful.

Although many cytosolic chaperones are highly abundant proteins thought to interact with a wide range of cytosolic proteins, only certain of these interacted with septins in our assay. If chaperones merely engaged the V_C_ portion of the septin-V_C_ fusions, and not the septins themselves, we might expect a much broader spectrum of cytosolic chaperone BiFC interactions. For example, the Hsp70 Ssa1 is the most abundant chaperone in the cytosol of unstressed yeast cells (Brownridge *et al*. 2013), but it did not interact with any septin at any temperature. Conversely, its much less abundant, 98%-identical paralog, Ssa2, interacted with all five septins (Table 4). Other studies using the same Ssa1-V_N_ strain were able to detect BiFC interactions with other proteins (Sung *et al*. 2013), demonstrating that the Ssa1-V_N_ fusion is functional. Ssa1 and Ssa2 also have distinct functions in prion propagation (Allen *et al*. 2005) and the vacuolar import and degradation pathway of gluconeogenic enzyme regulation (Brown *et al*. 2000; Sharma and Masison 2011). As another example, the Hsp70 Ssz1 forms a ribosome-associated complex with Zuo1 and the other Hsp70s Ssb1 or Ssb2, but Ssz1 appears to have lost its ability to bind nascent substrate polypeptides (Huang *et al*. 2005). In agreement with this idea, we found that Zuo1-V_N_ and Ssb1-V_N_, but not Ssz1- V_N_, interacted with V_C_-tagged septins (Table 4). Thus, we are confident that our BiFC results reveal specific associations, not non-specific interactions or promiscuous encounters.

A previous study using a trimolecular fluorescence complementation (TriFC) assay to identify septin-interacting proteins in living yeast cells specifically tested interactions with Hsp82 as a negative control, and found none (Finnigan *et al*. 2016). We suspect that the septin–Hsp82 interactions we see with BiFC reflect the relatively large sizes of the V_C_ and V_N_ tags, the linkers between them, and the C termini of the septins and chaperones to which they were fused, compared with the very small (20-21- residue) tags and linkers (18-33-residue) used in the TriFC system (Finnigan *et al*. 2016). The TriFC study demonstrated that the small tags and short linkers limit signal generation to interactions occurring within a short distance, such that only septins adjacent to each other in the septin hetero-oligomeric complex generate TriFC signal (Finnigan *et al*. 2016). In our BiFC system, all septins interact with all septins (our unpublished observations). Accordingly, Hsp82 could probably interact with any part of a septin protein and generate BiFC signal, but would likely need to bind nearby the N or C terminus in order to generate TriFC signal. Since our KADO model predicts that chaperones recognize septins with misfolded GTPase domains (Johnson *et al*. 2015), which are formed primarily by residues in the middle of septin proteins, using a split-fluorophore approach to identify septin–chaperone interactions may require larger tags and longer linkers.

Accordingly, the drop in specific chaperone BiFC signals we observed for mutant septins relative to their WT counterparts does not necessarily mean those chaperone– septin interactions no longer occurred. Differences in conformation or exposure of chaperone binding sites for a mutant septin may increase the physical distance between the C-terminal V_C_ tag on the septin and the C-terminal V_N_ tag on the chaperone beyond the BiFC threshold. These differences may explain diminished BiFC signals for chaperones whose deletions (Ydj1 and Gim3) perturb septin KADO, and whose overexpression (Ydj1) impedes mutant septin function.

The changes in chaperone interactions upon septin mutation were similar to changes for WT septins at high temperature (Table 4). Certain chaperones may recognize non-native conformations that WT proteins only sample at high temperature but mutant proteins readily adopt at low temperature. Recent work on mammalian Hsp90, for example, suggests that client binding depends on the combined probabilities of partial client unfolding, capture by Hsp70 of a transiently exposed site, and further unfolding events that expose adjacent client regions for capture by Hsp90 in complex with co-chaperones (Wang *et al*. 2020). The mutations we examined could promote any or all of these events. We chose V272M as a representative example of a p53 mutant with global structural instability and temperature-aggravated misfolding, and we have no reason to suspect that other p53 mutants with similar structural defects (manifested as temperature sensitivity, for example) would behave differently.

Alternatively, the presence of a mutant protein might itself alter chaperone expression in ways that mimic the effects of high temperature, such that a different repertoire of chaperones is available for interaction, independent of protein conformation *per se*. Indeed, expression of mutant p53 in yeast cells was sufficient to alter chaperone expression without an increase in temperature (Singh *et al*. 2010). The BiFC tags themselves may also alter chaperone expression: we detected BiFC signals with WT septins for chaperones whose expression in non-stress conditions is normally very low (e.g., Ssa3 (Werner-Washburne *et al*. 1987), Table 4). The V_C_ tag itself did not drive chaperone binding, however, since p53-V_C_ did not interact with Ssa3-V_N_ (Table 5).

While we think BiFC is a powerful tool for identifying chaperone interactions, chaperone binding assessed by any method does not indicate a functional role for that chaperone in protein folding. Whereas we do not yet know which chaperones are required for native septin folding, p53 folding has been more thoroughly studied and our BiFC interactions include chaperones that have not (yet) been implicated in p53 folding. Chaperones may “sample” a polypeptide during and after synthesis without changing the folding trajectory, and such transient sampling, irreversibly captured by BiFC, may explain at least some of the BiFC interactions we observed. Quantitative increases in chaperone BiFC signal for a mutant septin or p53 relative to WT could mean that otherwise inconsequential sampling by the chaperone of the WT protein has become consequential for the mutant, by altering its folding trajectory or occupying an oligomerization interface long/frequently enough to prevent native oligomerization. Chaperone binding at the septin–septin G interface and competition with other septins for interaction there is consistent with predictions from our KADO model for mutant septins.

By contrast, most known chaperone binding sites in WT p53 are found in the structurally labile DNA-binding domain (DBD) (Fourie *et al*. 1997; Müller *et al*. 2004), as opposed to the oligomerization domain. If the same sites mediate chaperone binding to p53(V272M), then it seems less likely that chaperone binding directly blocks oligomerization by mutant p53 and, since oligomerization buries the p53 nuclear export signal, indirectly promotes cytosolic sequestration. The three p53 nuclear localization signals are all located near the C terminus and outside the DBD (Shaulsky *et al*. 1990), hence direct occlusion of these sequences by chaperone binding is also an unlikely mechanism of cytosolic sequestration.

Instead, chaperone binding to the p53 DBD could inhibit nuclear localization of p53 in other ways. In mammalian cells, p53 nuclear localization is controlled by many factors not present in yeast, including the protein MDM2, intermediate filaments, and dynein-mediated transport along microtubules (reviewed in (O’Brate and Giannakakou 2003)). Nonetheless, the same NLS and NES sequences that control p53 nuclear localization in mammalian cells also control p53 nuclear localization when expressed in yeast (Godon *et al*. 2005). Binding of mammalian Hsp70–Hsp40 to the WT p53 DBD alters its conformation to disfavor DNA binding (Boysen *et al*. 2019), but an inability to bind DNA cannot explain the cytosolic sequestration of p53(V272M) because the contact mutant R273H does not have the same problems with nuclear retention (Figure 3B). We suspect that cytosolic chaperone–p53 complexes are retained in the cytosol because chaperone binding stabilizes a conformation of p53 in which the NLS is unavailable for proper interaction with the nuclear import machinery. Indeed, experiments in mammalian cells with a different TS p53 mutant, C135V, suggested that at high temperature binding of cytosolic Hsp70 chaperones to the mutant p53 renders the NLS inaccessible without directly occluding it (Akakura *et al*. 2001). p53 expressed in yeast can adopt several forms with distinct localization patterns – both punctate and diffuse, and in the cytosol, vacuole, or nucleus – some of which can be stably propagated in prion-like ways, suggesting conformational changes in p53 drive the changes in localization (Park *et al*. 2021). Since most of the chaperones we overexpressed are normally found diffusely in the cytosol, we think that the distinct patterns of p53 mislocalization outside the nucleus upon overexpression of distinct chaperones (punctate, diffuse, vacuolar) may have more to do with the conformation of p53 resulting from chaperone interaction than with chaperone binding *per se*. Along the same lines, vacuolar p53 upon Hsp90 inhibition probably represents p53 liberated from Hsp90 but captured by other chaperones and ultimately excluded from the nucleus.

Our findings that overexpression of any one of many distinct chaperones specifically inhibited the *function* of p53(V272M) suggest that cytosolic sequestration was sufficient but not necessary for functional inhibition. For example, Ydj1 overexpression inhibited p53(V272M) function but actually increased nuclear p53 levels (Figure 6, Figure S4A,B). Hence some conformations of p53 are presumably able to enter and remain in the nucleus but are unable to function there properly. From a broader perspective, these functional data with p53(V272M) add to our previous results with mutant septin function (Johnson *et al*. 2015) and run counter to the idea that elevated chaperone expression generally improves the folding and function of mutant proteins. While chaperone “assistance” with mutant p53 folding and function has been suggested in studies with other mutants expressed in yeast (including, interestingly, R273H) (Singh *et al*. 2010), our data demonstrate that overexpression of the same chaperone (Ssa2) was inhibitory to V272M function (Figure 6A). This discrepancy may point to differences in how chaperone binding alters the folding of a contact mutant (R273H) versus a conformational mutant (V272M). The extent and context of chaperone “overexpression” may also be important: the effect of Ssa2 on p53(R273H) was observed not by directly overexpressing Ssa2 but by adding ethanol, which indirectly induced Ssa2 (Singh *et al*. 2010).

Similarly, some of the chaperones that had a functional effect on p53(V272M) were negative for BiFC interaction (i.e., Ssa4, Hsp26, and Hsp31). One potential explanation for this discrepancy is the high *GAL1/10*-driven chaperone expression we used in the functional assays, compared to endogenous chaperone levels in BiFC. Finally, whereas the constitutively active Hsf1ΔN mutant inhibited the function of a mutant septin (Johnson *et al*. 2015), it had no effect on WT or mutant p53 (Figure 6A). Hsf1 drives expression of many different chaperones in the context of a balanced response to proteostatic stress (Morimoto 1998). While it may also indirectly upregulate other chaperones, high-level overexpression of a single chaperone perturbs this balance and likely drives p53 toward conformations that are only briefly/rarely sampled in the context of an Hsf1-driven stress response.

Another key point of comparison between our present results and those of prior studies is the effect of chaperone inhibition/deletion. In our earlier septin studies, deleting specific chaperone-encoding genes liberated mutant septins from apparent sequestration and allowed them to oligomerize with other, WT septins (Johnson *et al*. 2015). We focused on Ydj1 and Gim3 because published positive genetic interactions between *ydj1Δ* or *gim3Δ* and TS septin alleles hinted at improved function of the mutant septin when the chaperone was removed (Costanzo *et al*. 2010). In fact, however, the double-mutant cells were no more functional than either single mutant; they were merely less dysfunctional than one would have expected if each mutation independently compromised proliferation (Costanzo *et al*. 2010). Our finding that the Hsp90 inhibitor radicicol promotes nuclear localization of mutant p53 is conceptually similar, if we interpret it as p53 liberation from cytosolic chaperone sequestration but without fixing the folding problems that compromise function.

Indeed, our observation that radicicol also drives WT and R273H molecules of p53 into the nucleus but actually reduces overall p53 function (Figure 7) may explain the unexplained finding (Godon *et al*. 2005) that various forms of cellular stress (radiation, cadmium, and mercury) promote localization of p53 to the yeast nucleus. If these stressors (all of which are known to induce the heat shock response) generate misfolded or damaged proteins, those proteins may exceed the sequestration capacity of the cytosolic chaperone machinery, and liberate misfolded p53 molecules to enter the nucleus and interfere with the function of properly folded p53. Such a mechanism would be reminiscent of the way proteostatic stress liberates Hsf1 from cytosolic Hsp70– Hsp90 sequestration and allows Hsf1 to trimerize, accumulate in the nucleus and activate chaperone gene expression (Morimoto 1998). Not all ways of inhibiting chaperone sequestration have the same effect, however: deleting the gene encoding the small heat shock protein Hsp26 rescues the function of p53(R273H) expressed in yeast (Singh *et al*. 2010). Presumably, p53(R273H) is able to achieve a functional conformation without Hsp26 assistance, whereas Hsp90 contribution to p53 folding is indispensable. It follows that liberation of nonfunctional p53 molecules might have implications in chaperone inhibitor therapy. Chaperone upregulation in many tumors promotes the growth of cancerous cells by enabling “chaperone addicted” oncogenes or otherwise providing proteostatic benefit (Whitesell and Lindquist 2005). In tumor cells, chaperone binding to some p53 mutants protects the mutants from proteolysis (Walerych *et al*. 2004; Muller *et al*. 2008). However, if some liberated mutants escape degradation, inhibiting chaperones could permit a shift to dominant-negative p53 activity.

Finally, our findings do not support a KADO model for the recessive behavior of p53(V272M). KADO predicts delayed arrival of mutant V272M molecules in the nucleus and delayed mixing with WT, together with a recessive phenotype at all temperatures. Instead we observed a thermodynamic blockade on the ability of the vast majority of V272M molecules to reach a conformational state competent for nuclear localization. Sparing of WT inhibition by p53(V272M) and p53(V272M R273H) was only achieved at 37°, a temperature where the mutant was thermodynamically sequestered in the cytosol, rather than simply kinetically delayed in its nuclear arrival. For mutant septins, a poorly defined “window of opportunity” for hetero-oligomerization in the G1 phase of the cell cycle enforces mutant exclusion by KADO (Schaefer *et al*. 2016). We suspected that co-translational dimerization might represent a p53 “window of opportunity”, whereby a slow-folding mutant would be forced to dimerize post-translationally and miss the chance to form mutant dimers that could mix with WT dimers. However, based on our findings it is unclear if in yeast WT p53 dimerization is co-translational, and there is likely no kinetic “deadline” by which a p53 molecule must fold in order to properly oligomerize. Taken together, this study highlights key differences between septin oligomerization and p53 oligomerization and the effects of TS mutations and chaperone binding, which helps us understand the general principles underlying dominant and recessive mutant phenotypes.

## Acknowledgements

We thank Jeff Moore (University of Colorado Anschutz Medical Campus) for strains yJM3164 and yJM1838. We thank Chandra Tucker (University of Colorado Anschutz Medical Campus) for strain TH5; plasmids HSF1ΔN, Pdfr1-DH-p53-FR, and Pdfr1-DH-p53(Y220C)-FR; and access to the Yeast ORF collection. We thank Mark Johnston (University of Colorado Anschutz Medical Campus) for access to the Yeast Deletion collection. We thank Peter Kaiser (University of California Irvine School of Medicine) for strains RBy33 and RBy159. We thank Richard Iggo for plasmid pLS76. We thank Galit Lahav (Harvard Medical School) for plasmids p662, p663, p665, p666, p667, p668, p675, p678, and p679. We thank Bert Vogelstein for plasmids pCMV-p53(R249S) (Addgene plasmid #16438; http://n2t.net/addgene:16438; RRID:Addgene_16438) and RS314-H (Addgene plasmid #17541; http://n2t.net/addgene:17541; RRID:Addgene_17541).

## Funding and additional information

This work was supported by the National Institute of General Medical Sciences of the National Institutes of Health under Award 5R01GM124024 (to M.A.M.) and Award T32GM008730 (to A.S.D), by the Alzheimer’s Association under award NIRGD-12-241119, and by the Paul R. Ohara II Seed Grant Program and University of Colorado Cancer Center ACS-IRG Grant. The content is solely the responsibility of the authors and does not necessarily represent the official views of the National Institutes of Health.

## Abbreviations and nomenclature

BiFC: bimolecular fluorescence complementation
DHFR: dihydrofolate reductase
KADO: kinetic allele discrimination in oligomerization
TS: temperature sensitive
WT: wild type
YFP: yellow fluorescent protein

## Figure legends

**Figure S1.**
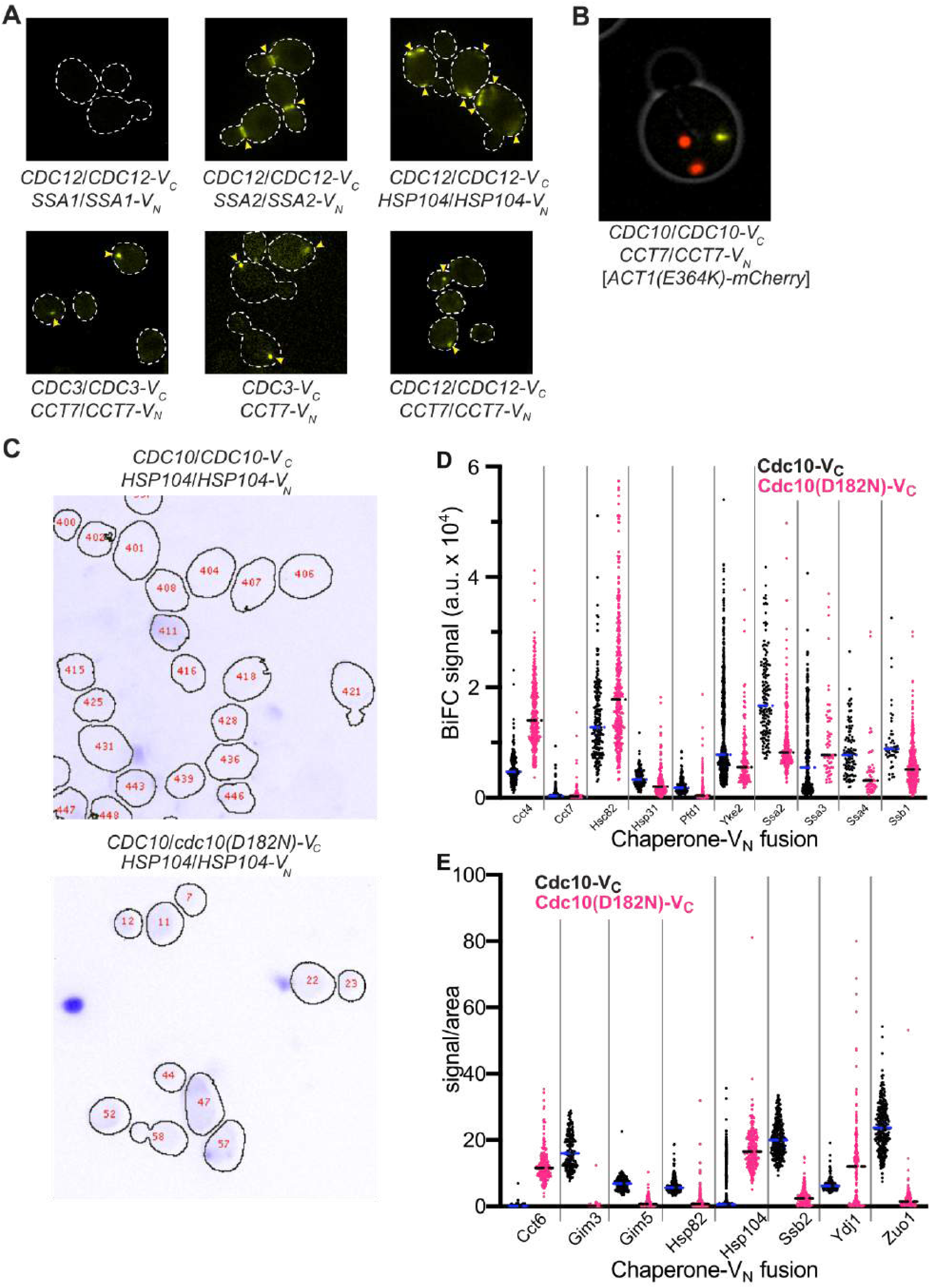
Characterization of BiFC signals generated from septin–chaperone interactions. (A) Log-phase cultures of cells of the indicated genotypes were imaged for transmitted light and YFP fluorescence. The transmitted light images were used to generate cell outlines (dashed white lines). Yellow arrowheads indicate discrete BiFC signals. “*CDC3-V_C_ CCT7-V_N_*” indicates the haploid strain H00526. Other strains were diploids made by mating the haploid strains H00383, H00384, H00387, H00377, YO802 and YO685 in appropriate combinations. (B) Diploid cells co-expressing Cdc10-V_C_ and Cct7-V_N_ made by mating strains H00383 and YO1057 and carrying a plasmid encoding Act1(E364K)-mCherry under control of the *GAL1/10* promoter (pDK304) were cultured in galactose-containing medium to drive Act1(E364K)-mCherry overexpression and shifted to 37° for 2 hours, to drive Act1(E364K)-mCherry aggregation, prior to imaging. Shown is a representative overlay image of the transmitted light, YFP, and mCherry signals. (C) Representative images showing cell outlines overlaid on transmitted images (inverted and false-colored blue) used for measuring intracellular fluorescence. (D) Cellular BiFC signal away from bud necks was quantified for diploid cells co-expressing Cdc10-V_C_ or Cdc10(D182N)-V_C_ and the indicated V_N_-tagged chaperones. The number of cells analyzed for each genotype ranged from 46 to 460. Strains were diploids made by mating YO1057 or H06530 with H00381, H00383, H00399, H00397, H00393, H00389, H00388, H00403, H00404, or H00395. (E) As in (C) but for the indicated chaperones and the values plotted represent the ratio of signal to area for each cell. Strains were diploids made by mating YO1057 or H06530 with H00382, H00392, H00390, H00398, H00384, H00394, H00385, or H00396.

**Figure S2.**
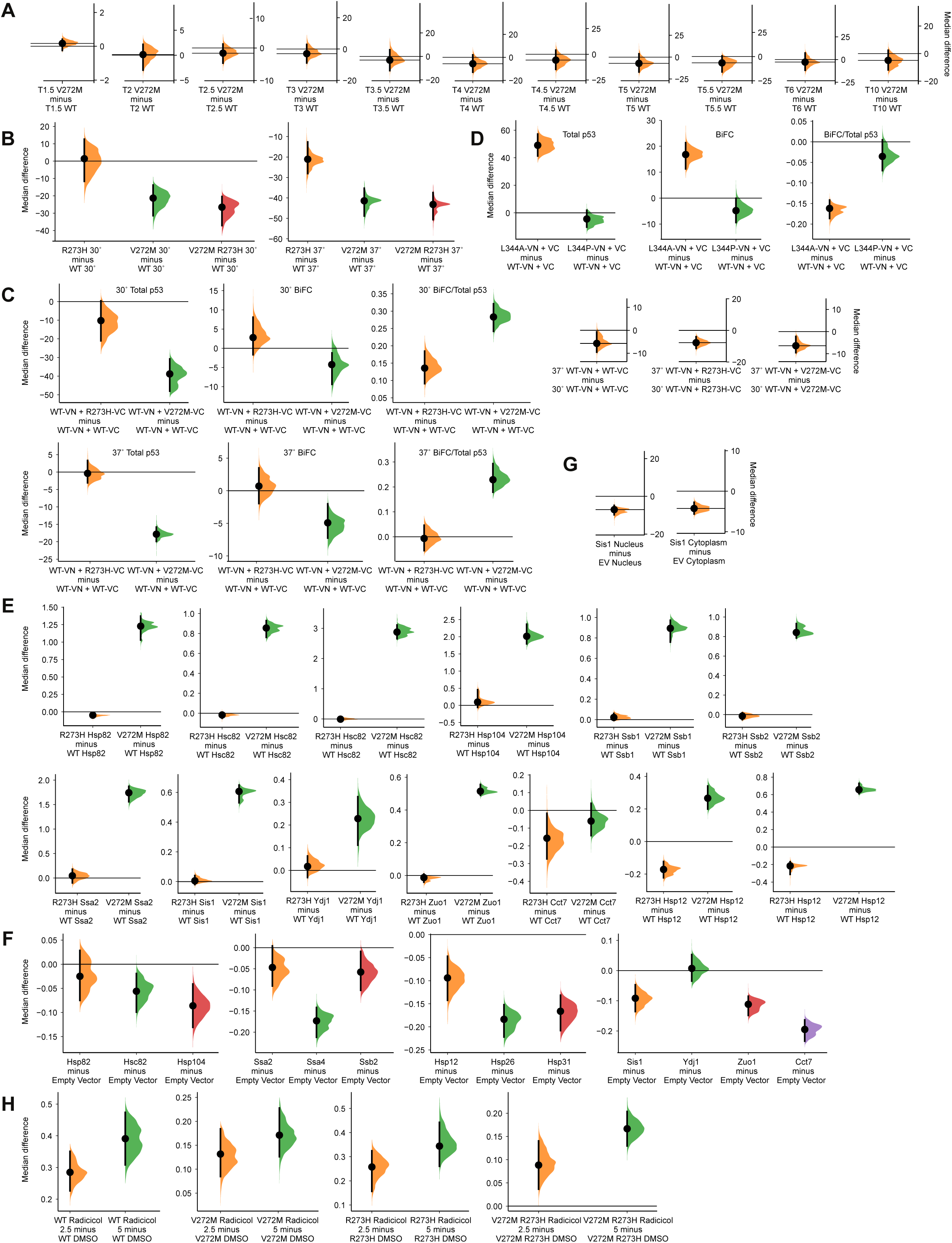
Effect size estimations of changes in p53 fluorescence signals. (A) For the data in Figure 3A, the median difference for nuclear mKate2 comparisons between p53(V272M) and WT p53 are shown in Cumming estimation plots as bootstrap sampling distributions. Each median difference is depicted as a dot. Each 95% confidence interval is indicated by the ends of the vertical error bars. (B) For the data in Figure 3B, the median difference for nuclear mKate2 comparisons between the indicated p53 mutants and WT p53 are shown in Cumming estimation plots as bootstrap sampling distributions. Each median difference is depicted as a dot. Each 95% confidence interval is indicated by the ends of the vertical error bars. (C) For the data in Figure 3C, the median difference for nuclear total p53 (mKate2) comparisons, nuclear BiFC (Venus) comparisons, and nuclear BiFC/total p53 comparisons between the indicated co-expressed BiFC alleles are shown in Cumming estimation plots as bootstrap sampling distributions. Each median difference is depicted as a dot. Each 95% confidence interval is indicated by the ends of the vertical error bars. (D) For the data in Figure 4B, the median difference for nuclear total p53 (mKate2) comparisons, nuclear BiFC (Venus) comparisons, and nuclear BiFC/total p53 comparisons between the indicated co-expressed BiFC alleles are shown in Cumming estimation plots as bootstrap sampling distributions. Each median difference is depicted as a dot. Each 95% confidence interval is indicated by the ends of the vertical error bars. (E) For the data in Figure 5B, the median difference for BiFC/total p53 comparisons between p53(R273H) or p53(V272) and WT p53 are shown in Cumming estimation plots as bootstrap sampling distributions. Each median difference is depicted as a dot. Each 95% confidence interval is indicated by the ends of the vertical error bars. (F) For the data in Figure 6C, the median difference for nuclear/cytosolic mKate2 comparisons between chaperone-overexpressing cells and empty vector are shown in Cumming estimation plots as bootstrap sampling distributions. Each median difference is depicted as a dot. Each 95% confidence interval is indicated by the ends of the vertical error bars. (G) For the data in Figure 6D, the median difference for nuclear and cytosolic mKate2 comparisons between Sis1-overexpressing cells and empty vector are shown in Cumming estimation plots as bootstrap sampling distributions. Each median difference is depicted as a dot. Each 95% confidence interval is indicated by the ends of the vertical error bars. (H) For the data in Figure 7A, the median difference for nuclear/cytosolic mKate2 comparisons between radicicol- and DMSO-treated cells are shown in Cumming estimation plots as bootstrap sampling distributions. Each median difference is depicted as a dot. Each 95% confidence interval is indicated by the ends of the vertical error bars.

**Figure S3.**
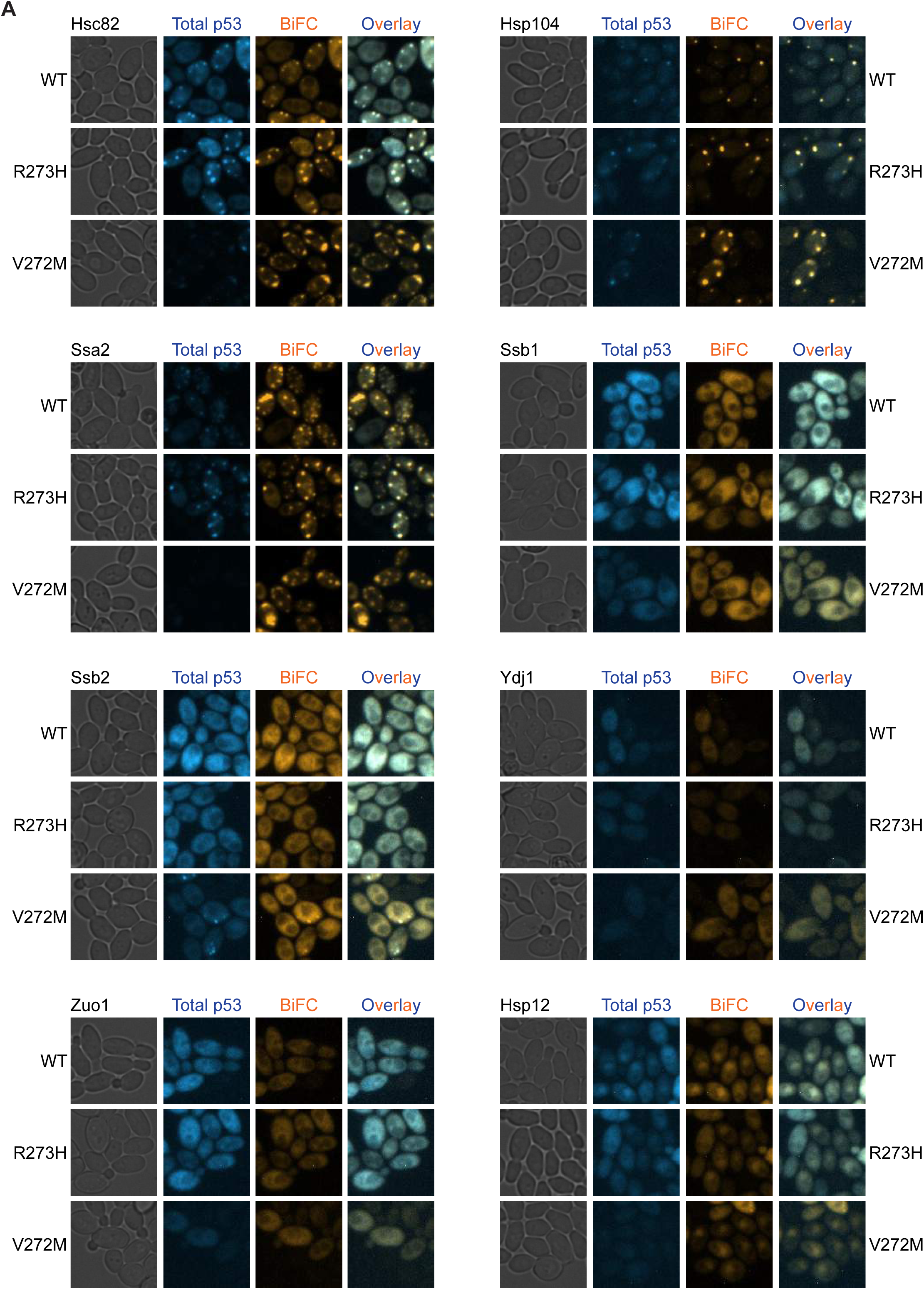
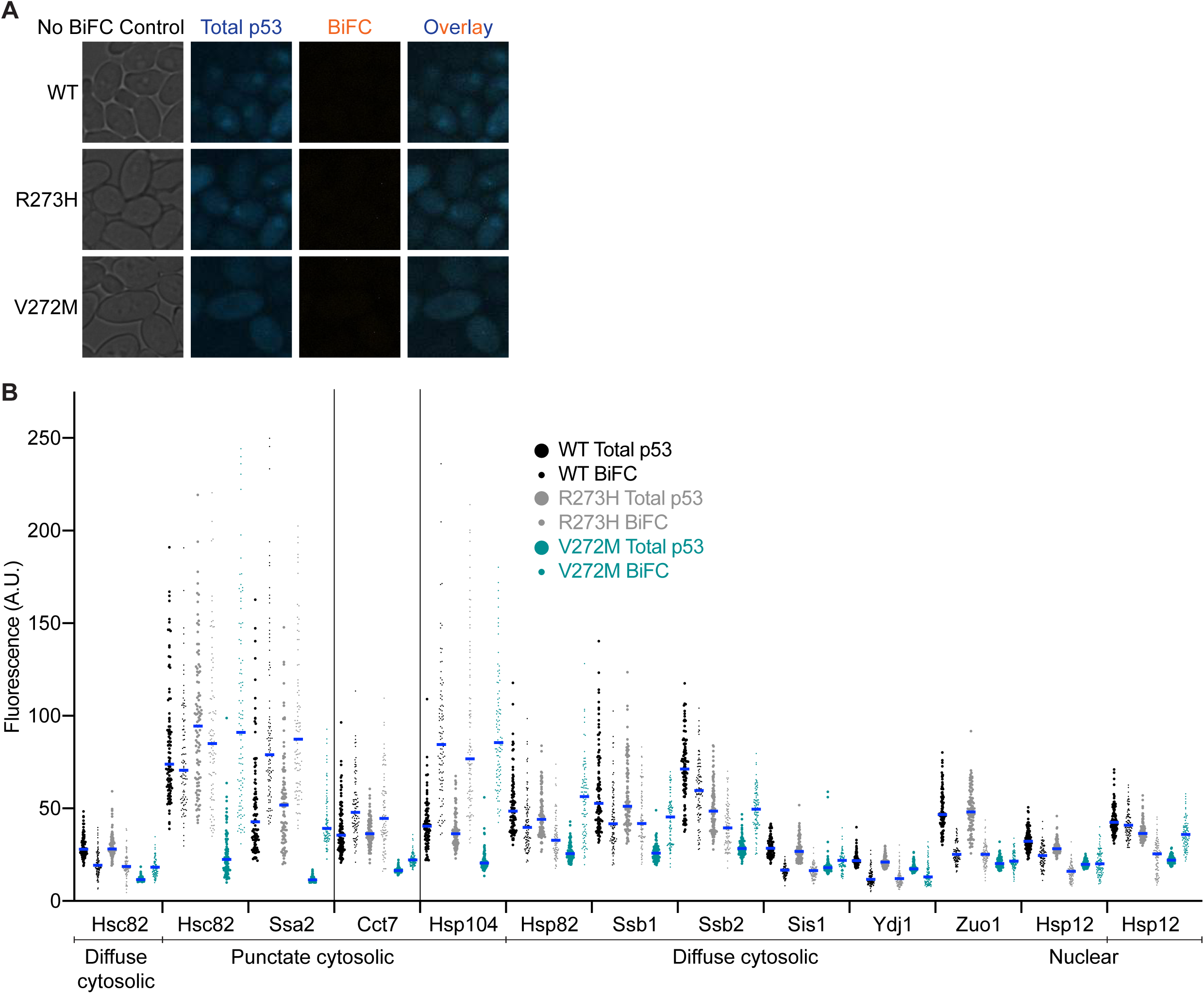
Extended characterization of cytosolic chaperone-p53 interactions. (A) Representative micrographs of p53–chaperone BiFC interactions in cells expressing Hsc82-V_N_, Hsp104-V_N_, Ssa2-V_N_, Ssb1-V_N_, Ssb2-V_N_, Ydj1-V_N_, Zuo1-V_N_, Hsp12-V_N_, or in cells expressing empty vector (“No BiFC Control”), and carrying plasmids encoding the indicated p53-mKate2-V_C_ alleles. Total p53 (mKate2) fluorescence images were false-colored blue and BiFC (Venus) fluorescence images were false-colored orange. Prior to overlay, brightness and contrast were adjusted equivalently for each set of images. Plasmids were pMAM110, “WT”, pMAM112, “R273H”; pMAM111, “V272M”; pRS316 (not labeled). Strains were diploids made by mating BY4742 cells carrying the p53- mKate2-V_C_ plasmids with H00399, “Hsc82”; H00384, “Hsp104”; H00388, “Ssa2”; H00395, “Ssb1”; H00394, “Ssb2”; H00385, “Ydj1”; H00396, “Zuo1”; or H06684, “Hsp12”. “No BiFC Control” was a diploid made by mating BY4742 cells carrying the p53-mKate2-V_C_ plasmids with BY4741 cells carrying pRS316. (B) Total p53 (mKate2) and BiFC (Venus) fluorescence in cells cultured at 30° carrying plasmids encoding the indicated p53-mKate2-V_C_ alleles together with the indicated V_N_-tagged chaperone. Blue lines denote median values. Vertical lines group the samples imaged with the same settings (LED intensity and exposure time). The subcellular location where the BiFC interaction was quantified is indicated below. Plasmids were pMAM110, “WT”, pMAM112, “R273H”; pMAM111, “V272M”. Strains were diploids made by mating BY4742 cells carrying the plasmids with H00399, “Hsc82”; H00388, “Ssa2”; H00383, “Cct7”; H00384, “Hsp104”; H00398, “Hsp82”; H00395, “Ssb1”; H00394, “Ssb2”; H06680, “Sis1”; H00385, “Ydj1”; H00396, “Zuo1”; or H06684, “Hsp12”.

**Figure S4.**
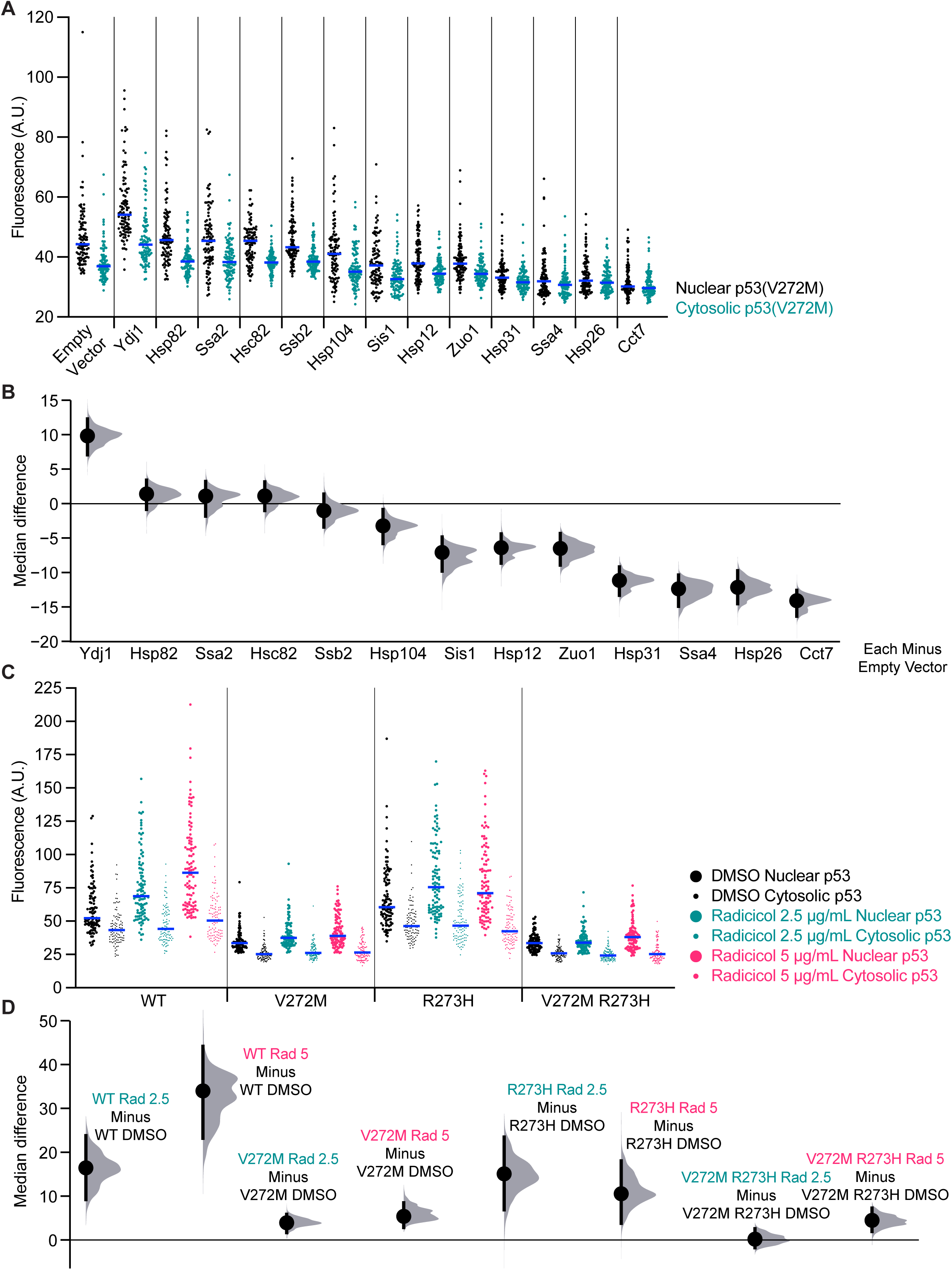
Raw nuclear and cytosolic p53 levels in chaperone-overexpressing cells and Hsp90-inhibited cells. (A) Nuclear and cytosolic mKate2 fluorescence in diploid cells carrying a plasmid encoding p53(V272M)-mKate2-Venus together with the indicated *GAL1/10*-driven chaperone plasmid or an empty vector. Cells were grown to mid-log phase at 30° with galactose to overexpress chaperones prior to imaging. Blue lines denote median values. Vertical lines separate genotypes. Plasmids were pMAM86, “p53(V272M)”; pRS424, “Empty Vector”; G00353, “Ydj1”; G00630, “Hsp82”; G00633, “Ssa2”; G00799, “Hsc82”; G00634, “Ssb2”; G00631, “Hsp104”; G00632, “Sis1”; G00627, “Hsp12”; G00635, “Zuo1”; G00629, “Hsp31”; G00789, “Ssa4”; G00628, “Hsp26”; G00620, “Cct7”. Strains were diploids made by mating yJM3164 cells carrying pMAM86 with yJM1838 cells carrying the indicated *GAL1/10*-driven chaperone plasmid or empty vector. (B) For the data in (A), the median difference for nuclear mKate2 comparisons between chaperone-overexpressing cells and empty vector are shown in Cumming estimation plots as bootstrap sampling distributions. Each median difference is depicted as a dot. Each 95% confidence interval is indicated by the ends of the vertical error bars. (C) Nuclear and cytosolic mKate2 fluorescence in yJM3164 cells carrying plasmids encoding the indicated alleles of p53-mKate2-Venus. Cells were grown to mid-log phase at 37° with the indicated concentration of radicicol or equivalent volume of DMSO prior to imaging. Blue lines denote median values. Vertical lines separate genotypes. Plasmids were pMAM85, “WT”; pMAM86, “V272M”; pMAM87, “R273H”; pMAM104, “V272M R273H”. (D) For the data in (C), the median difference for nuclear mKate2 comparisons between radicicol- and DMSO-treated cells are shown in Cumming estimation plots as bootstrap sampling distributions. Each median difference is depicted as a dot. Each 95% confidence interval is indicated by the ends of the vertical error bars.

